# Increased recognition memory precision with decreased neural discrimination

**DOI:** 10.1101/2024.06.19.599765

**Authors:** Jeremie Güsten, David Berron, Gabriel Ziegler, Emrah Düzel

**Affiliations:** German Center for Neurodegenerative Diseases, Magdeburg, Germany; Institute of Cognitive Neurology and Dementia Research, Otto-von-Guericke University, Magdeburg, Germany; Institute of Cognitive Neuroscience, University College London, London, United Kingdom

**Keywords:** Mnemonic Discrimination, Training-induced Plasticity, Cognitive Training, task fMRI

## Abstract

Mnemonic discrimination (MD), the ability to distinguish between similar experiences in memory, is essential for memory precision. We hypothesized that training mnemonic discrimination should improve memory precision by increasing the neural activity differences between a new experience and a similar familiar experience (repeat). Participants performed a 2-week, web-based training program. For group 1, memory similarity was adapted to performance. Group 2 trained non-adaptively and group 3 served as active control. Adaptivity improved training gain and led to transfer to other memory tasks. Surprisingly, training reduced neural activity differences between new (lure) and similar familiar (repeat) experiences. Thus, instead of enhancing neural activity to lures and decreasing neural activity to repeats, training led to increased activity to repeats. In several brain regions this pattern was associated with improved MD. These findings highlight an unexpected neural mechanism underpinning improved memory precision in recognition memory.

## Introduction

Mnemonic discrimination (MD), the ability to distinguish between similar experiences, possibly relies on efficient hippocampal pattern separation (PS), the orthogonalization of neural representations. The contributions of the hippocampus to MD have been shown in rodent (Gilbert et al., 2001) and human research (Brock Kirwan et al., 2012; Reagh & Yassa, 2014). In human fMRI, lure discrimination (LD), the usually positive difference between activation to lure vs. repeats, is often used as a proxy for PS and is typically linked to hippocampal subfields DG/CA3 (Bakker et al., 2008; Berron et al., 2016; Lacy et al., 2011). Recent views suggest that PS may actually occur throughout cortex (Aimone et al., 2011; Kent et al., 2016; Rolls, 2016), and LD signals have been reported in extra-hippocampal MTL (Reagh & Yassa, 2014) as well as the ventral visual stream (Chouinard et al., 2008; Klippenstein et al., 2020; Koutstaal et al., 2001; Nash et al., 2021; Pidgeon & Morcom, 2016; Wais et al., 2017). The importance of ventral visual regions, especially perirhinal cortex, in the MD of objects has long been known (Bussey et al., 2003), consistent with the hierarchical- representational account according to which object representations become more complex and hence less overlapping along the ventral-visual stream (Kent et al., 2016). However, object disambiguation is not necessarily linked to positive LD (Lure > Repeat), since repetition enhancement has also been reported along the visual stream (Chouinard et al., 2008; Klippenstein et al., 2020; Koutstaal et al., 2001; Nash et al., 2021; Pidgeon & Morcom, 2016; Wais et al., 2017). Also, it is unclear whether ventral-visual LD is directly linked to MD. For instance, Klippenstein et al. observed LD in occipital cortex, but found no correlation with discrimination performance (MD). Apart from ventral-visual regions, the prefrontal cortex has been shown to exhibit LD (Pidgeon & Morcom, 2016) and has been linked to MD in rodents (Johnson et al., 2021). While most research has focused on object stimuli, it is not known to what extent the neocortex also supports MD of scenes or spatial stimuli, given that objects/scene stimuli are subserved by the distinct ventral/dorsal visual streams, respectively (Ranganath & Ritchey, 2012; Rolls, 2023). Increasing evidence suggests that separation may already occur before reaching HC in dorsal stream areas such as medial entorhinal cortex (Reagh & Yassa, 2014) and PHC (Aly et al., 2013; Reagh & Yassa, 2014).

To date, it is unclear whether MD performance can be improved by cognitive training. This question is relevant because impairment of MD is seen with advancing age and with Alzheimer’s disease pathology (Maass et al., 2019; Berron et al., 2018; Fidalgo et al., 2016; Güsten et al., 2021; Holden et al., 2013; Reagh et al., 2016; Stark et al., 2013). Studies have shown improvements in MD performance following mild physical exercise (Suwabe et al., 2018) or playing visually rich computer games (Clemenson et al., 2020; Clemenson & Stark, 2015). However, the mechanisms inducing these changes remain unclear given that brain activation patterns elicited by the training task were not measured, and no training-induced effect on standard neuropsychological tests was observed (Clemenson et al., 2020; Clemenson & Stark, 2015). An associated question is whether any cognitive training related improvement of MD would also lead to transfer to other memory tasks.

An important consideration in the design of cognitive trainings is the role of adaptivity. Lövdén et al., 2010 propose that change in ability is caused by a prolonged mismatch between the current ability and demands posed by a task, stressing the importance of adaptivity in inducing ability improvements and potential transfer to untrained tasks. However, in the field of working memory training results are mixed, with some finding a benefit of adaptivity (Brehmer et al., 2012; Flegal et al., 2019; Holmes et al., 2009), while others do not (Hotton et al., 2018; Ripp et al., 2022). Whether these findings are transferable to MD training is unclear, and the only training studies testing effects on MD were non-adaptive (Clemenson et al., 2020; Clemenson & Stark, 2015). Moreover, while in n-back working memory trainings difficulty is modulated by the span of stimuli interposed between a presentation and test stimulus (Jaeggi et al., 2008; Colom et al., 2013; Pahor et al., 2022), MD training should aim to decrease the discriminability of stimuli in order to tax PS more strongly.

Given that the neural LD response is seen as a physiological hallmark of MD, we hypothesized that training MD should improve memory precision by increasing LD, reflecting better neural separation of lures and repeats. That is, the neural activity differences between lures and repeats should increase with MD training and this increase should be correlated with behavioral improvement in MD performance. To test this hypothesis, young adults performed individualized training on a modified version of the well-studied (Berron et al., 2018, 2019) object-scene MD task using a custom-made web-based training platform (MemTrain). In study 1, we examined whether an adaptive task-regime benefits 1) task performance increase and 2) near transfer to other memory tasks. To target PS more directly, stimulus discriminability was reduced instead of increasing the memory (n-back) span. Study 2 included task-fMRI, allowing us to assess MTL regions displaying LD at baseline and to subsequently investigate training-induced LD changes in MTL and across neocortex. Given distinct cortical streams, we distinguished between domain-specific (objects versus scenes) and domain-general patterns of change. In order to link activation change with behavior, we correlated ΔLD and ΔMD in regions showing mean change and 2) explored regions showing training-specific correlations between ΔLD and ΔMD. Finally, we assessed training-related near and far transfer to a range of cognitive tasks, including a task of associative recall of objects placed within three dimensional scenes (Object-in-Scene, OiS, task) using very similarly designed stimuli as in the MD training. This allowed us to test whether improving neural discrimination would also benefit neural association.

## Results

### Adaptiveness of training leads to greater training improvement across domains in object-scene task

The main aim of study 1 was to assess the effect of adaptiveness on training outcome. We explored this question using Pr as a performance measure. Crucially, a group x session interaction for object performance (F(2, 52) = 3.7863, p = 0.02916) and a similar trend for scene performance (F(2, 51.5) = 2.1894, p = 0.1223) were observed (fig. 1A). Groupwise contrasts for objects showed a stronger increase for the adaptive (t(52) = 5.450, p < .0001) than the non-adaptive training (t(52) = 2.417, p = 0.0192) and the control group, which showed no significant improvement at all (0.05, t(52) = 1.480, p = 0.1450). In the scene condition, the pattern of improvement was similar (adaptive: t(51) = 4.805, p < .0001; non-adaptive: t(51) = 2.642, p = 0.0109; control: t(51.9) = 1.592, p = 0.1174). We then tested, whether the training effect also differed between the two training groups. And indeed, we found differences in training improvement for the two groups regarding object performance (F(1, 36) = 4.3942, p = 0.04315) and a trend for scene performance (F(1, 36) = 2.5754, p = 0.1173). Given that results for objects were more robust, we directly explored whether performance improvement differed as a function of the stimulus domain. This is quite reasonable, given that object and scene stimuli partially rely on different brain networks (Berron et al., 2018; Ranganath et al., 2012), which may in turn differentially react to training. We thus set up an LME including session, group and domain as well as all their higher-order interactions. The session x group x domain interaction was not significant however (F(2, 155) = 0.3804, p = 0.6842), suggesting that the training effect was similar across domains, while the session x group effect persisted (F(2, 155) = 6.5184, p = 0.0019). Moreover, the group x session effect for scenes was in fact significant (F(1, 35) = 5.0491, p = 0.0311) if removing clear outliers (mean absolute deviations > 2.5, N = 2). We finally investigated how hit rates or false alarm rates contributed to the performance improvement. Across domains, training-induced change was more strongly linked to FAR (objects: F(2, 52) = 7.8930, p = 0.001; scenes: F(2, 52) = 2.3505, p = 0.1054), then hit-rates (objects: F(2, 52) = 0.1753, p = 0.8397; scenes: F(2, 52) = 1.8106, p = 0.1737). While in study 2 a more difficult 6-back version of the task was used, and no non-adaptive training group was included, the pattern of results was very similar (fig. 1B). That is, we observed stronger, domain-independent performance improvements for the training group mainly linked to a decrease in FAR. Given that MD improvement was linked to FAR reductions, we henceforth use FAR as a proxy for MD.

**Figure 1.**
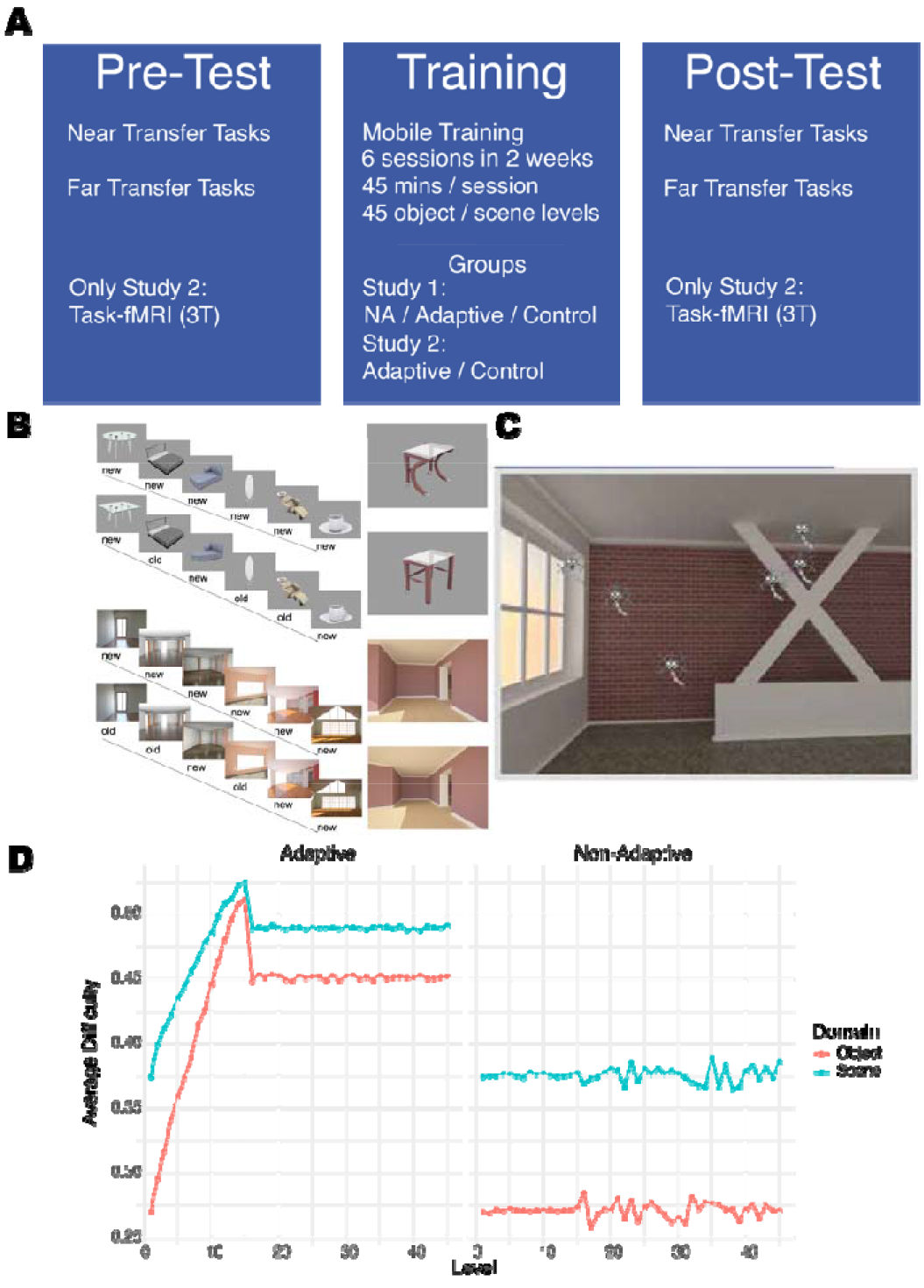
Overview of the training design. A. Participants underwent a computerized training, from a remote but quiet location (mobile). Pre- and Post-tests were on near and far transfer tasks, with a 2 weeks training period in between. Both training and control groups underwent active training, but on different tasks. **B** The training group performed a 6 -back version of the object-scene task (Berron et al., 2018, Güsten et al., 2021), with separate object and scene trials, and domain-specific level increase. In the adaptive training, level difficulty was increased by showing more difficult images based on previously evaluated difficulty. **C** The control group training consisted of eliminating moving neurons by clicking on them. With level increase, neurons became more numerous and faster. To reduce stimulus differences between tasks, background images were equivalent to the scenes from the training task. **D Average level difficulty of the training task.** Stimulus discriminability rather than trial length was modified to more directly target discrimination ability rather than long-term memory. In the adaptive group, level difficulty increased almost linearly, and plateaued at high difficulty in later levels, while in the non-adaptive group it remained constant at a low level. Average stimulus difficulty was estimated from an independent study (Güsten, 2021) and was measured across trial types (lure vs. repeat trial) and stimulus version order. Due to stimulus availability constraints, scene trials were generally more difficult than object trials.

### Training-related improvements in Object-in-Scene transfer task require adaptivity

We next investigated, whether training affected overall performance increases in near transfer tasks, starting with the Object-in-Scene (OiS) task. Including adaptive, non-adaptive and control group, we did observe a significant group x session interaction regarding the number of correct responses (F(2, 106) = 4.56, p = 0.013, see fig. 1C). Groupwise contrasts revealed that the adaptive group improved more between sessions (*t(*106) = 6.048, p < .001), while change was not significant for the control group (*t(*106) = 1.968, p = 0.052) or the non-adaptive group (*t(*106) = 1.089, p = 0.279). As there are two error types in the task (internal and external), we explored whether improvements were uniquely associated to one. However, for both error types the adaptive group showed improvement (external: *t(*106) = 1.811, p = 0.073; internal: *t(*106) = 2.493, p = 0.014), while the non-adaptive (external: *t(*106) = 0.958, p = 0.34; internal: *t(*106) = 0.466, p = 0.643) and control group (external: *t(*106) = 0.63, p = 0.53; internal: *t(*106) = 1.428, p = 0.156) did not. Importantly, effect sizes were also much larger in the adaptive compared to the non-adaptive group, which cannot be explained by the unequal sample size (N_adaptive_=46, N_non-adaptive_=19). As for the MST, we observed no group x session interaction, suggesting no substantial effect of the training on performance in this task.

Given a lack of transfer effects to the near-transfer MST task, we did not expect to find any effects in far transfer tasks. However, we observed a group x session interaction in the delayed recall of the verbal learning task, as indicated by a group x session interaction effect (F(1, 48) = 6.334, p = 0.0152). The pairwise contrasts show a negative slope for controls (*t(*48) = -2.1455, p = 0.037), and a positive slope for the training group (*t(*48) = 1.3861, p = 0.1721), reflecting performance increase in delayed recall for the controls and a non-significant decrease for the training group. It has to be noted however, that baseline performance was very different for the two groups, where in the pre-session the training group actually remembered more items at delayed than immediate recall (see supp. fig. 1). Finally, no training effects were observed in the remaining transfer measures (supp. tables 5- 7).

### Perirhinal cortex and parahippocampal cortex show lure discrimination profile at baseline

We next explored functional activation during task performance before training across the MTL to identify regions showing LD, indicative of neural lure-repeat discrimination. First, repetition sensitivity was assessed across *a-priori* selected MTL ROIs at baseline, since this effect is hypothesized to underlie the LD effect. As expected, all ROIs showed a pattern of repetition suppression (First > hit; see fig. 2). Furthermore, no ROI showed a domain-specific difference in the amount of RS, suggesting that information pertaining to both domains is processed in each of these MTL regions. Having established repetition sensitivity, we went on to test LD across ROIs, and found effects in perirhinal cortex (PrC) (*t(*160)=-5.436, p<0.001, p_fdr_<0.001), parahippocampal cortex (PhC) and observed (*t(*160)= -2.838, p=0.005, p_fdr_=0.018) and an effect in CA3, which remained a trend after fdr-correction (*t(*156)= 2.171, p=0.031, p_fdr_=0.073). Thus, while both PrC and PhC showed a familiar pattern of positive LD, CA3 showed the opposite pattern of greater activation for hits than CRs (negative LD). Moreover, we found no group-level effect of LD in any other ROI. In summary, only PrC and PhC showed a pattern consistent with novelty signaling during correct lure trials, suggesting a potential role in MD.

**Figure 2.**
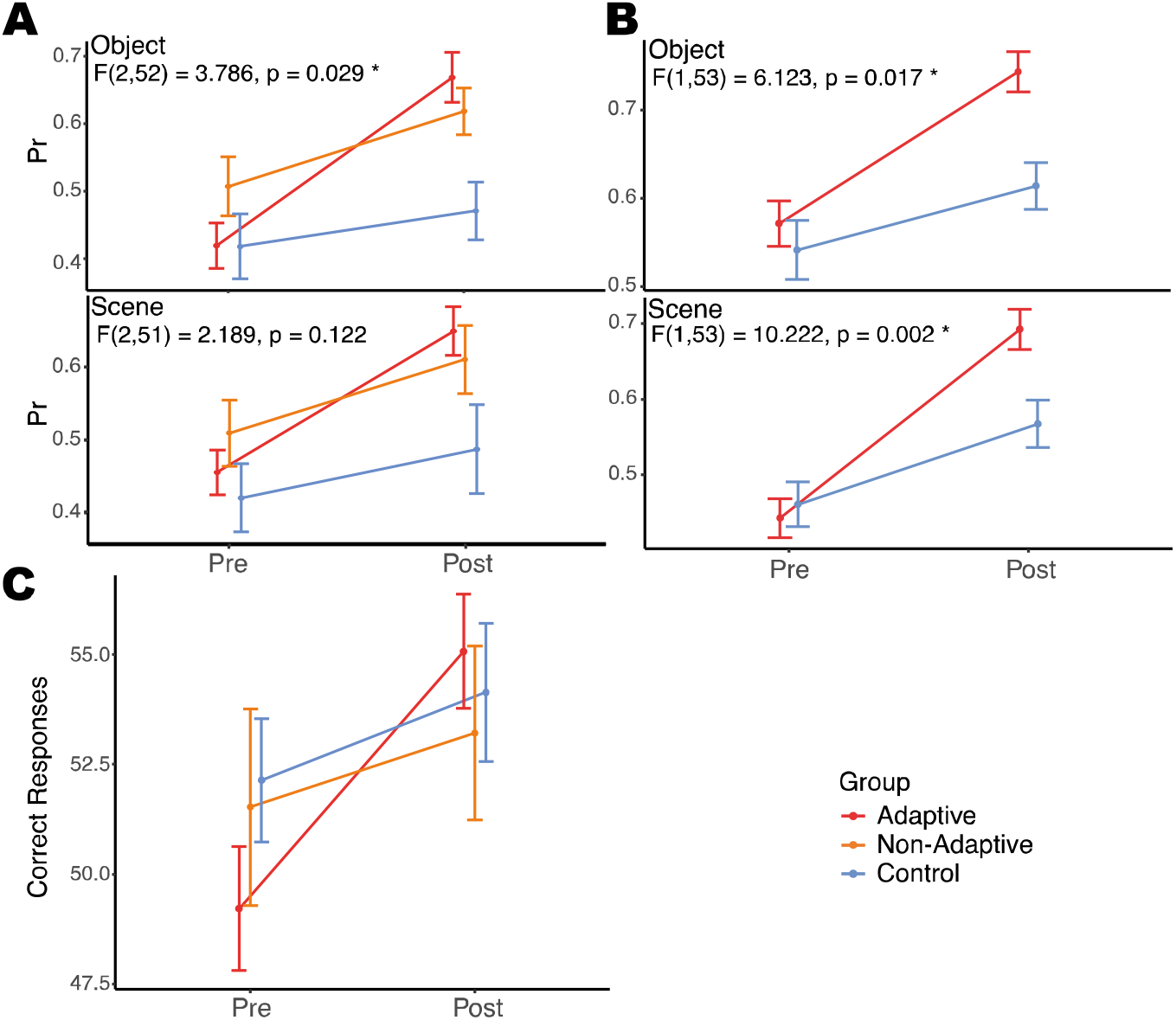
Cognitive training improves mnemonic discrimination. (**A**) Study 1: 2- back object-scene task performance (Pr), by group and session. Object as well as scene discrimination performance, the adaptive training group shows the steepest training- related improvement. Significant group x session interaction effect for objects, and similar trend for scenes. Figure shows mean Pr and standard error (SE). (**B**) Study 2: 6 - back object-scene task performance, by group and session. As in the original 2 -back task in study 1 (**A**), the training group shows greater task improvements than controls, in both object and scene discrimination. (**C**) Object-in-Scene (OiS) transfer task performance across both studies. The training group shows a steeper increase in correct cued object-in-scene recall, while improvements are only marginal (control group) or non-significant (non-adaptive group).

### Cognitive training leads to decrease of object-related lure discrimination in perirhinal cortex

To determine the impact of training on LD enhancement, which we hypothesized may underlie the observed MD improvements, we investigated two scenarios: one involving a domain-general change, whereby a region displays increased lure-repeat discrimination across stimulus types; and the other, a domain-specific change indicative of training-induced refinement in discriminating a particular stimulus type. No domain-general change was observed, however. On the other hand, when looking at domain-specific change, PrC showed a significant group x session x domain effect (*t(*153)=2.814, p=0.004, *p_fdr_*= 0.026). Two-way models for each domain separately showed that the group x session effect in PrC was significant for objects (*t*(51)=-2.612, *p*=0.012), but not scenes (*t*(51)=1.661, *p*=0.103). Finally, pairwise comparisons revealed the effect for objects to be related to a decrease of LD in the training group (*t*(26)=-2.11, *p*=0.045), without significant change in controls (*t*(25)=1.57, *p*=0.13). As can be seen in fig. 3, this training-related decrease in LD was mainly due to an increase in activity for hits. Finally, the voxelwise whole-brain analysis showed no clusters of group x session or group x session x domain effect on LD. Thus unexpectedly, training led to no LD increase in any region, and even induced object-specific LD decrease in PrC.

**Figure 3.**
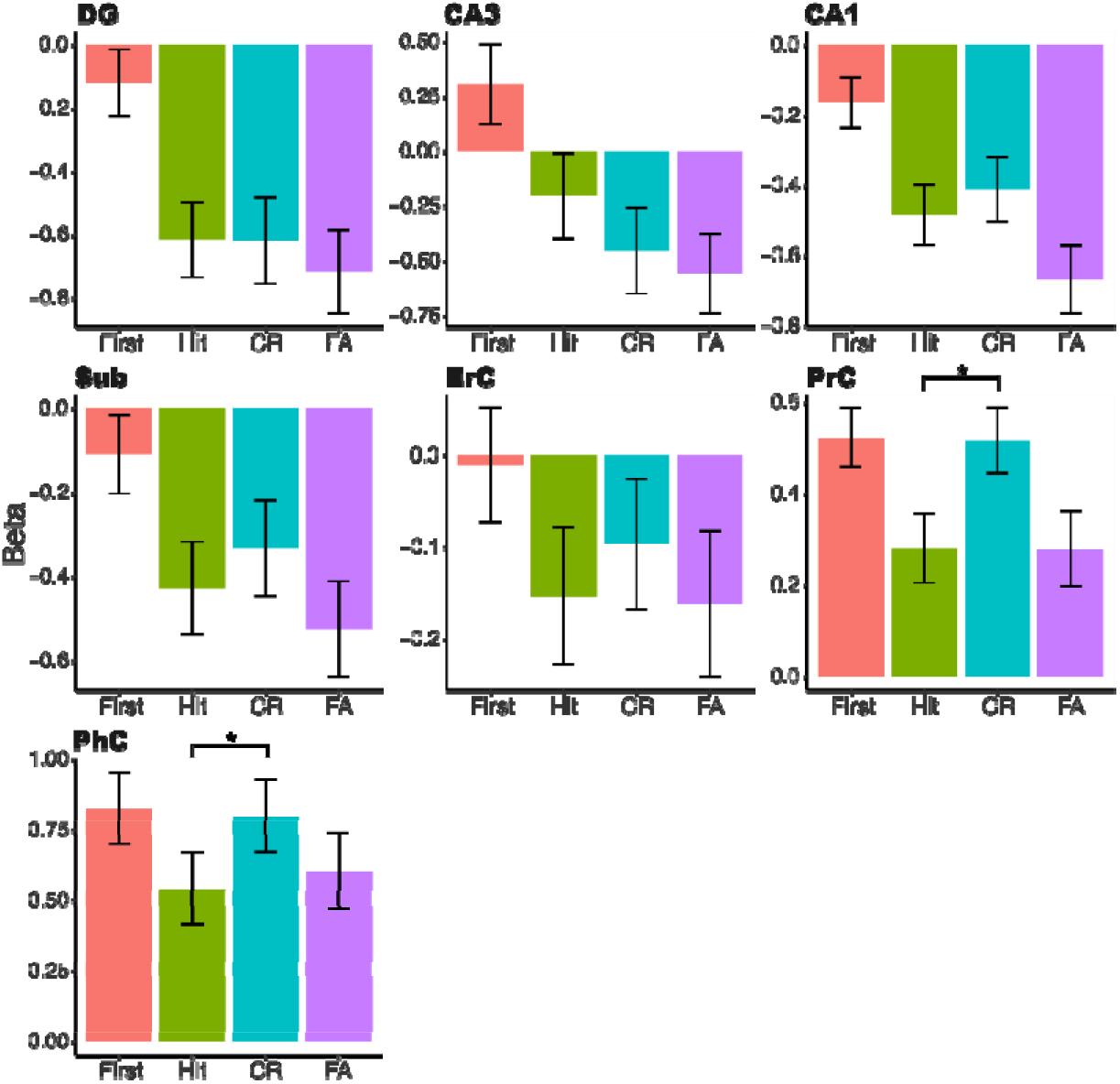
Activation profile at baseline across MTL subfield regions. Figure shows condition-specific beta estimates for first presentations (red), hits (green), correct rejections (blue) and false alarms (purple). Scrambled trials were used as implicit baseline. PrC and PhC show lure discrimination (LD) profile (significant difference between hits and CRs), linked to a lure novelty response. CA3 showed a marginally significant effect of LD, linked to a weaker response to correct lures than to hits. Effects are fdr-correct at alpha = 0.05.

### CA3, CA1 and left medial frontal cortex show increased correct retrieval-related activity after training

Beyond examining changes in LD, we assessed an orthogonal effect: the average activation during correct retrieval trials (hits + correct rejections). A change in this measure might reflect altered neural efficiency, with decreases suggesting enhanced efficiency or reduced task reliance, and increases pointing to heightened engagement. Again we tested both domain-general and domain-specific change across MTL ROIs, and found significant domain-general effects in CA3 (*t*(155)=3.924, *p*<0.001, *p_fdr_*<0.001), Sub (*t*(153.4)=3.153, *p*=0.002, *p_fdr_*=0.007), CA1 (*t*(156.2)= 2.333, *p*=0.021, *p_fdr_*=0.049). Pairwise comparisons revealed that across these ROIs correct retrieval activation increased in the training group (CA3: *t*(26)=2.662, *p*=0.013; CA1: *t*(25)=2.34, *p*=0.028; Sub: *t*(25)=2.03, *p*=0.053) without significant change in controls (CA3: *t*(24)= -1.006, *p*=0.325; CA1: *t*(25)= 0.027, *p*=0.979; Sub: *t*(24)=-0.858, *p*=0.399). Unlike for LD, no domain-specific training effect was observed. Similar to the hippocampal effects, a voxelwise whole-brain analysis revealed a cluster of domain-general correct retrieval increase spreading across the left medial frontal and medial superior frontal cortex (see fig. 4B), but no domain-specific effect. A follow-up simple slope analysis of the cluster showed that the interaction was linked to an increase in correct retrieval activity in the training group (*t*(96)= 3.327, *p*=0.001), with no significant change in the controls (*t*(96)= -0.884, *p*=0.183).

**Figure 4.**
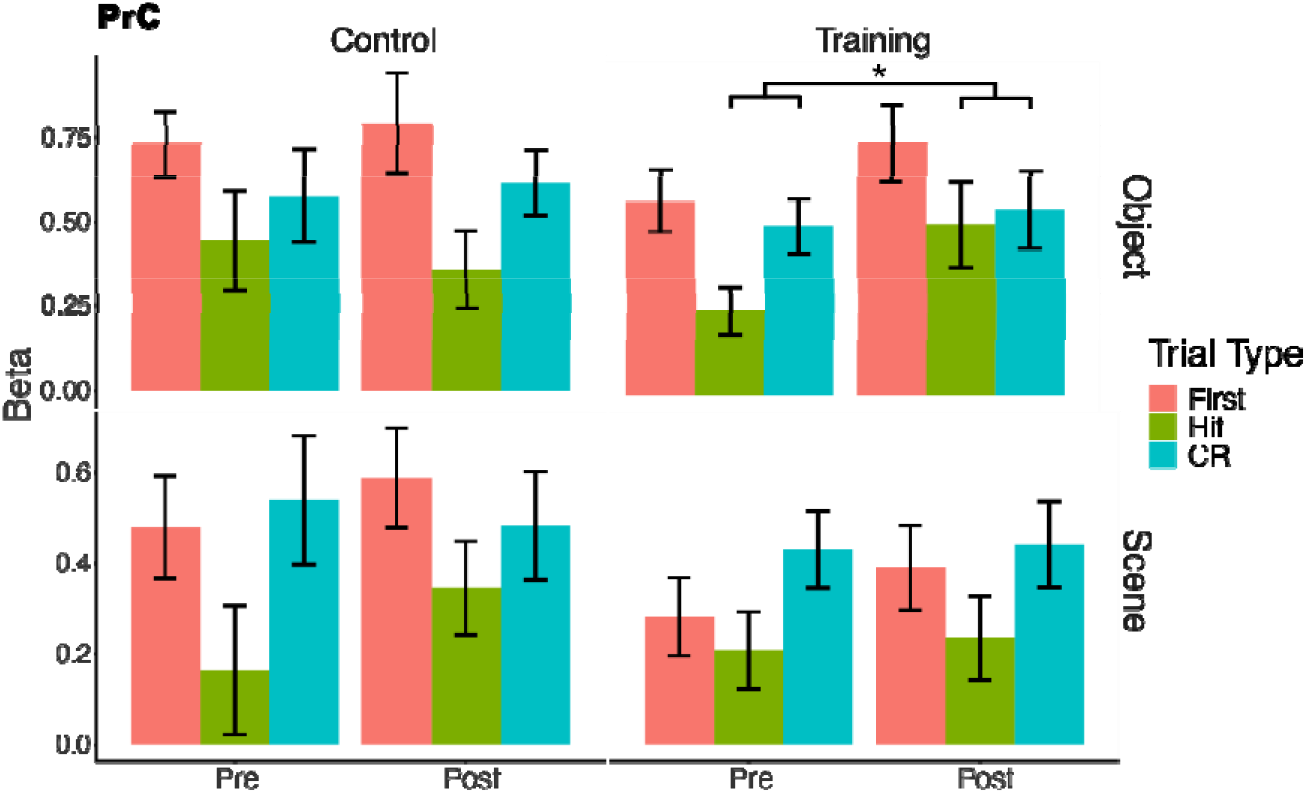
PrC shows object-specific change of LD in training group. Figure shows beta estimates for first presentations, hits and correct rejection, separated by session and group and domain. A domain-specific training effect on LD was found in PrC, shown by significant group x session x domain interaction. A significant group x session effect for objects was linked to LD change in the training group. While CR activity remained similar in this group, the effect is driven by an increase in hit activity. * Shows significant session effect on LD in follow-up pairwise comparison for object trials in training group, at alpha = 0.05 (uncorrected).

### Absence of correlation between false alarm rate changes and activation changes in regions showing training-induced activation shifts

We then investigated whether the observed activation changes were associated with MD improvements in the training group. However, no significant correlation was found between object-specific ΔLD activation and ΔFAR in PrC (*F*(1, 25)= 0.866, *p*=0.395). Moreover, investigating the link between Δ(correct retrieval) activation in CA3 and CA1 with ΔFAR revealed no effect for CA3 (*t*(25)= 0.544, *p*=0.591). Nonetheless, we observed a unexpected since positive trend for CA1 (*t*(24)= 1.737, *p*=0.095), linking increases in correct retrieval activation to less reductions in FAR. In addition, follow-up analyses on the extracted left medial frontal cluster showed no significant associations between Δ(correct retrieval) and ΔFAR. In summary, the results suggest that the observed training-induced activation shifts show no linear association with MD improvements.

### Training-induced associations between lure discrimination decrease and mnemonic discrimination improvements in domain-general and object-specific clusters

Given the lack of a linear association between activation changes and improvements in MD in regions showing training-induced activation shifts, we expanded our analysis to explore other regions’ involvement in the observed MD enhancements. Thus, we examined how training influenced the relationship between FAR and activation changes, using an activation change x group interaction model. Starting with LD, a domain-general cluster was found in supramarginal gyrus (figure 6; table 1), linked to positive ΔLD - ΔFAR correlation in the training group (*t*(96)= 3.616, *p*<0.001), but no correlation in controls (*t*(96)= 0.041, *p*=0.967). Interestingly, the region showed preferential activation to CR at baseline, with no differences between first and hit activation (at baseline; see supp. fig. 2), suggesting a lack of repetition suppression. Follow-up analysis showed that the interaction was significant for hit activation separately (*t*(96)= -2.98, *p*=0.004), linked again to significant correlation in the training group (*t*(96)= -3.878, *p*<0.001) and no effect in controls (*t*(96)= -0.15, *p*=0.881), but not for CR activation. Thus the observed effect was mainly linked to training-induced Δ(hit activity) - ΔFAR correlation.

**Figure 5.**
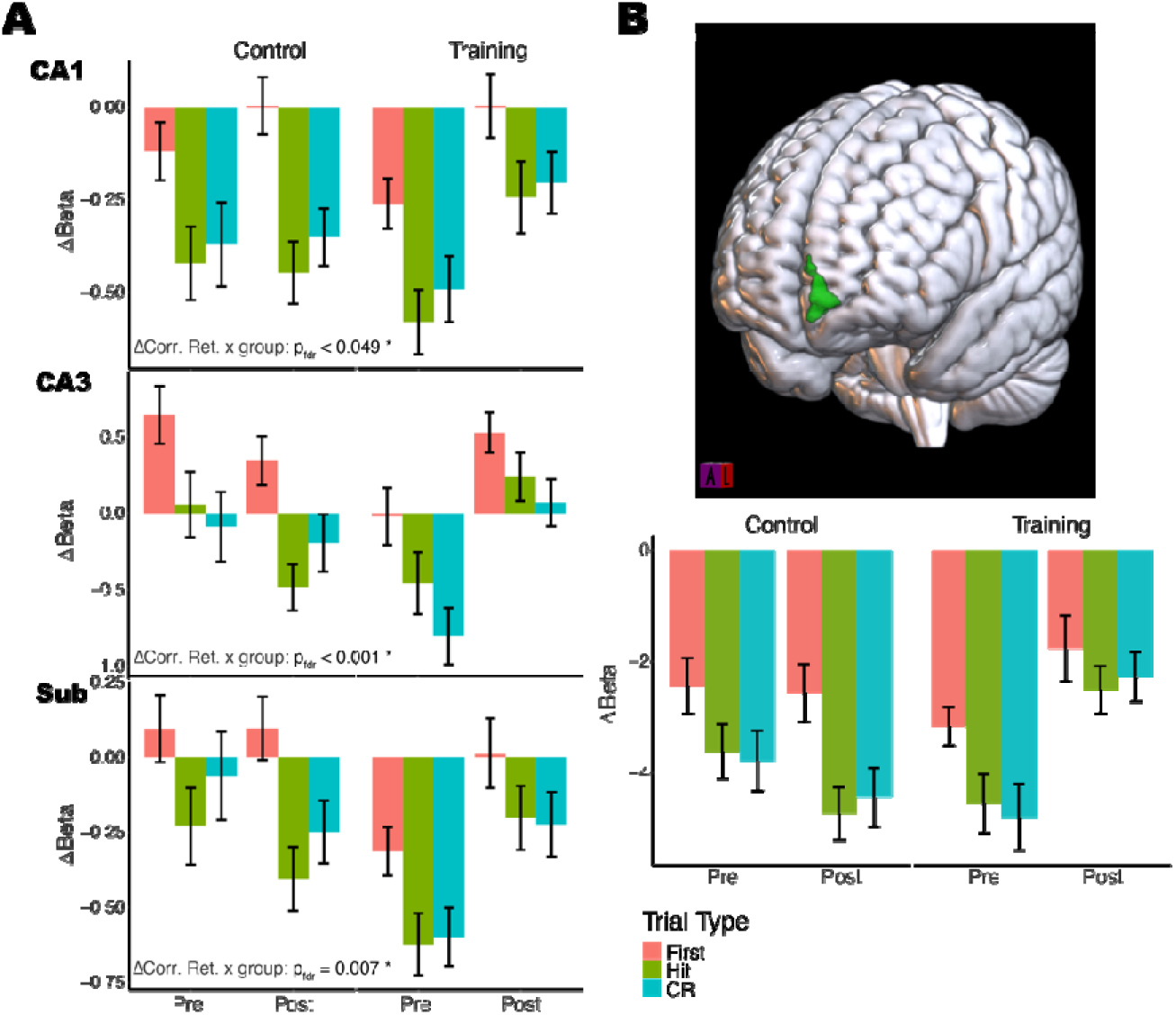
Training-related change in correct retrieval activity. (**A**) Condition- specific activations show training-related increases in correct retrieval activity (hits + correct rejections) in three MTL regions: CA1, CA3 and Sub. All three regions show an increase in activity only for the training group. (**B**) An exploratory whole-brain analysis revealed a cluster of training-specific correct retrieval activity increase in left medial and superior frontal cortex (shown in green). Significant cluster-level effect at alpha = 0.05, after FWE-correction and applying cluster-forming threshold at p=0.001.

**Figure 6.**
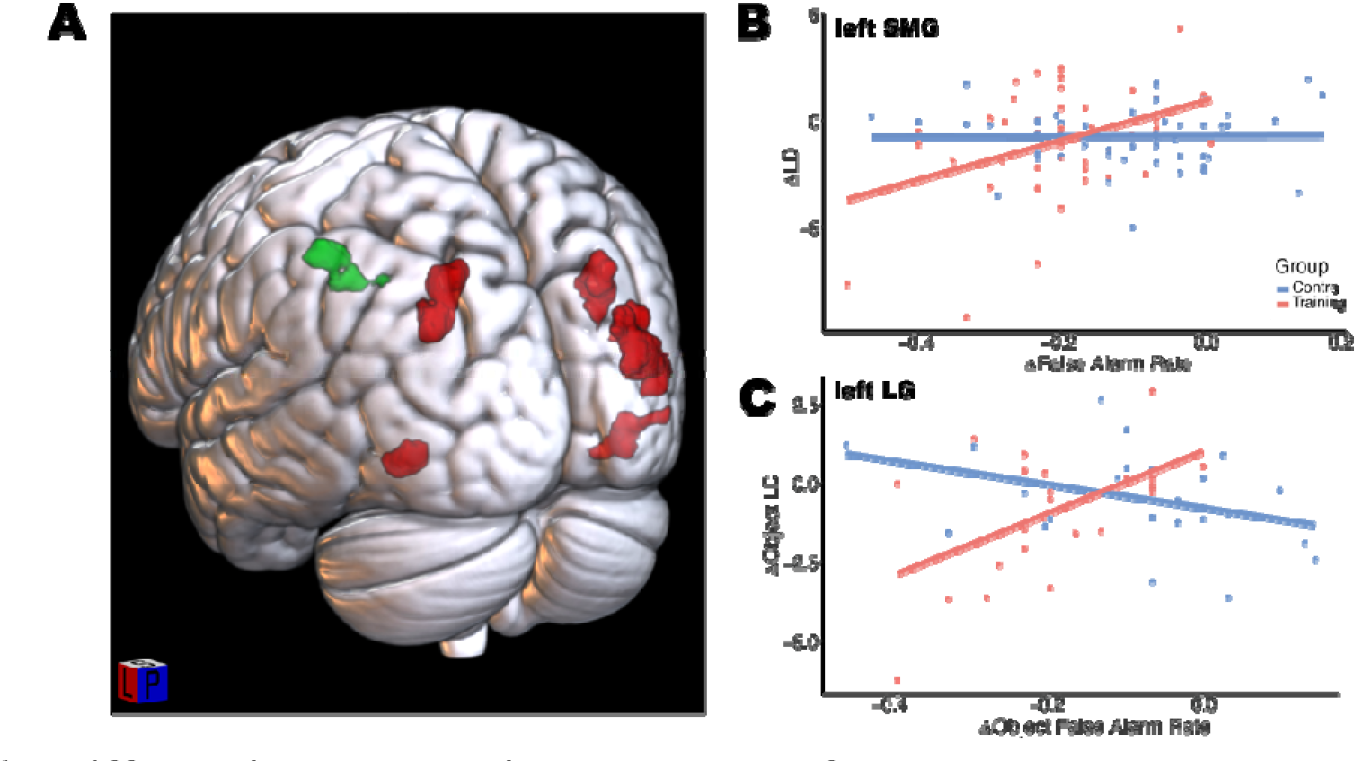
Differential correlations between false alarm rate change and lure discrimination change. (**A**) A whole-brain analysis of training-dependent Δobject CR activity - Δobject FAR correlation. It revealed a domain-general cluster in left supramarginal gyrus (shown in green), and object-specific clusters in right middle occipital gyrus, left lingual gyrus, left superior parietal lobe, right angular gyrus and right inferior temporal gyrus (all shown in red). For both domain-general (**B**) and object-specific effects (**C,** exemplified by lLG), the interaction was driven by a positive Δobject CR activity - Δobject FAR association in the training, but not the control group.

**Table 1.** Δ **(FAR) x group interaction effect on** Δ **(LD).** SMG=Supramarginal Gyrus, MOG=Middle Occipital Gyrus, SPL=Superior parietal lobule, AG=Angular Gyrus, ITG=Inferior Temporal Gyrus, LG=Lingual Gyrus

| Domain | Cluster | Peak | MNI (mm) |  |  |  | Region |  |
| --- | --- | --- | --- | --- | --- | --- | --- | --- |
|  | p(FWE-corr) | size | Z | x | y | z |  |  |
| **General** | 0.01 | 590 | 4.29 | -45 | -41 | 48 | R | SMG |
|  |  |  | 3.73 | -35 | -41 | 41 |  |  |
|  |  |  | 3.45 | -28 | -47 | 40 |  |  |
| **Object** | < 0.001 | 2035 | 4.9 | 37 | -86 | 19 | R | MOG |
|  |  |  | 4.2 | 42 | -79 | 9 |  |  |
|  |  |  | 3.77 | 36 | -79 | 29 |  |  |
|  | < 0.001 | 1594 | 4.27 | -23 | -71 | 43 | L | SPL |
|  |  |  | 3.9 | -24 | -67 | 29 |  |  |
|  |  |  | 3.71 | -31 | -73 | 30 |  |  |
|  | < 0.001 | 978 | 3.86 | 32 | -68 | 39 | R | AG |
|  |  |  | 3.63 | 28 | -69 | 26 |  |  |
|  | 0.013 | 556 | 3.87 | 45 | -61 | -7 | R | ITG |
|  |  |  | 3.44 | 42 | -53 | -16 |  |  |
|  |  |  | 3.43 | 51 | -72 | -4 |  |  |
|  | 0.019 | 516 | 4.32 | -23 | -57 | -8 | L | LG |

Additionally, domain-specific analyses pinpointed several visual and parietal regions where increased object-related activity correlated with better MD in the training group, but not in controls (figure 6; table 1). Interestingly, all clusters exhibited an association between object LD decrease and FAR reductions in the training group. As was the case for the domain- general cluster, the effect seemed to be linked primarily to the Δ(object hit activity) - Δ(object FAR) correlation across the clusters, with no significant correlation for Δ(object CR activity) -Δ(object FAR). Notably, no effect was detected for the change in scene LD.

In contrast to the observations for LD, our analysis revealed neither domain-general nor domain-specific alterations in the association between change in correct retrieval and FAR. These finding suggest a stronger link between MD enhancements and changes in LD rather than correct retrieval activation.

### Object-scene task performance and Object-in-Scene performance show similar association with object lure discrimination in overlapping regions

Lastly, we explored whether the observed training-induced functional changes were associated with OiS performance improvements. Starting with the pattern of LD change in PrC, we found no correlation between Δ(Sum Correct) and Δ(LD activity) in the training group, however. Further analyses across MTL ROIs revealed no significant training-related correlation change. In summary, as was the case for MD, OiS improvements showed no linear association with the observed LD changes in PrC.

Next, we correlated Δ(LD activity) with Δ(Sum Correct) in clusters having shown correlation change with FARs in the whole-brain analysis (domain-general: supramarginal gyrus; object-specific: occipital, parietal and inferior temporal clusters). While no effect was observed in the supramarginal cluster for either performance measure, we observed a significant negative correlation of Δ(Sum Correct) on Δ(object LD activity) in rITG (*t*(46)= -2.202, *p*=0.033), rAG (*t*(46)= -2.61, *p*=0.012), lSPL (*t*(46)= (-2.319, *p*= 0.025, see fig. 6C), and a trend in lLG (*t*(46)= -1.713, *p*=0.093). This suggests that across these clusters, individual decreases in object LD were associated with increase in correct OiS responses in the training group in addition to MD improvements. Moreover, the link was not significant in any of these regions for the control group. When testing the interaction directly, no region reached a significant Δ(Sum Correct) x group effect, however.

### Correct retrieval activation shows negative training-induced association with Object-in-Scene performance across several medial temporal regions

We investigated correlations between correct retrieval activation and OiS performance changes (fig. 6), by first examining MTL regions showing mean activation changes, namely, CA1, CA3 and Subiculum. However, only CA3 showed a trend for a negative link between Δ(Correct retrieval) and Δ(Sum Correct) in the training group (t(23)= -1.945, p=0.064). In addition, a significant Δ(Sum Correct) x group effect (t(45)= -2.215, p=0.032) showed that this link was specific to the training group (fig. 6). Finally, we asked whether correlation change occurred in other MTL areas. Indeed, we found a Δ(Sum Correct) x group effect in PhC (t(47)= -4.209, pfdr<0.001). While Δ(correct retrieval) was positively linked to Δ(Sum Correct) in the control group (t(47)= 3.499, p=0.001), the reverse was found in the training group (t(47)= -2.58, p=0.013). Further Δ(Sum Correct) x group were found in ErC (t(4)= - 2.388, pfdr=0.074) and Subiculum (t(45)= -2.031, pfdr=0.084). The pattern was similar to PhC, with a positive link in controls (ErC: t(45)= 2.306, p=0.026; Sub: t(45)= 1.856, p=0.070) and non-significant negative links in the training group (ErC: t(45)= -1.211, p=0.232; Sub: t(45)= -1.097, p=0.278). This finding is surprising, as it suggests that correct retrieval activation increases, as were observed in the hippocampus, were detrimental to transfer task improvements. In summary, the relationship between training-induced functional changes and OiS task enhancements seems to be complex, with some adaptations seemingly benefitting improvements, while others may hamper them.

## Discussion

In two studies, we assessed the effect of adaptive vs. non-adaptive training on a 6-back MD task over two weeks. Adaptive training enhanced performance more than non-adaptive training, as evidenced by scores on the trained task, a 2-back version of the task as well as positive transfer to an Object-in-Scene (OiS) task, though not to another MD task (MST). Baseline activation in the MTL showed very robust lure discrimination (LD) in PrC and PhC, while training led to a significant decrease of object LD in PrC. Moreover, several hippocampal regions (CA1, CA3, Sub) and the medial frontal cortex showed increased activation during correct retrieval after training. Yet these changes did not correlate with individual FAR decreases. However, domain-general LD decrease in supramarginal gyrus, and object-specific LD decrease in visual and parietal areas was associated with FAR reductions and OiS performance improvements in the training group.

The fact that adaptivity was necessary to induce performance improvements on a near transfer task echoes findings from working memory training (Brehmer et al., 2011; Flegal et al., 2019) and theoretical work (Lövdén et al., 2010). Using adaptive and non-adaptive training also served as a control for simple task familiarity. However, our design did have certain limitations: first, contrary to the control task, difficulty did not increase throughout all levels due to limited stimulus availability, effectively removing adaptivity at higher levels (see fig. 8D). But if anything, full adaptivity should have increased the differences found. Moreover, level progression did not accelerate beyond the difficulty plateau (sup. Fig. 4B), and perceived difficulty did not differ between training group and controls (see feedback questionnaire in supplementary), suggesting that the task remained similarly challenging.

Lack of transfer to the only MD transfer task -the MST- in combination with transfer to the more distant (in terms of a distinction between discrimination and association) OiS task suggests a benefit unrelated to hippocampal pattern separation, which should be a shared physiological mechanisms by our task and the MST. Also, transfer to OiS performance was likely not driven by increased HC activation given the negative training-induced correlation between hippocampal Δ(correct retrieval) and Δ(OiS performance). While both the MST and object-scene task target pattern separation, they differ considerably in terms of list length and stimulus pair similarity: the object-scene task uses very subtle stimulus pair differences, whereas the MST tests for the detection of broader, within-category change. It has been argued that pattern separation happens at different levels of ambiguity (Kent et al., 2016), and our participants may have learned to resolve mnemonic ambiguity at a lower representational level than is required for the MST. Accordingly, all regions showing a link between LD change and performance increase were located in visual as well as posterior parietal cortex, which has been associated with the retrieval of mnemonic detail (Cabeza et al., 2008; Hutchinson et al., 2009; Ramanan et al., 2018). To clarify this, future studies should include MD transfer tasks requiring resolution of lower-level ambiguity. On the other hand, we found transfer to OiS performance, and both OiS and object-scene task improvements were associated with object LD decrease in overlapping regions. Changes in object processing may thus have facilitated improvements on both training and transfer task. As a limitation, it should be noted that certain scenes used in the OiS and object-scene task showed similarities, but the fact that improvement was mainly linked to object processing suggests that this was not the main driver of transfer.

We had hypothesized that regions exhibiting LD at baseline, in this case the PrC and PhC, would show increased LD after training. Suprisingly however, training led to decreased object LD in PrC. PrC had shown the clearest LD profile at baseline (fig. 4), adding to previous fMRI studies showing robust object LD in that region (Reagh & Yassa, 2014). Interestingly, the object-specificity of LD change likely reflects the region’s bias towards objects processing (supp. fig. 3). The domain-specific decrease in LD and the fact that PrC has long been linked to visual object discrimination (Barense et al., 2010; Clarke & Tyler, 2014; Lee et al., 2006; McLelland et al., 2014) suggests that its role in object retrieval changed with training. Moreover, we found no evidence of LD increase or decrease outside of the MTL. However, training induced a link between Δ(FAR) and Δ(object LD) in occipital, parietal and inferior temporal regions, whereby LD change became negatively associated with MD change through training. These findings complement the current literature by showing that LD signals outside the MTL can become associated with performance, contrary to previous cross-sectional data (e.g. Klippenstein, 2020). LD decrease mostly reflected a weakening of lure novelty rather than increased repetition enhancement, given that all regions showing the effect exhibited a lure novelty profile at baseline (supp. fig. 2). Interestingly, MD improvements in these regions were particularly linked to increased activation to hits, while LD change in PrC was also driven by changes in hit activation. More generally, the results suggest that improvements in object MD relied on changes in the processing of repeats, mainly in object-specific regions of the ventral-visual stream.

There may be several explanations for a training-induced decrease in LD. On the one hand, participants may have learned to pay attention to relevant item features. It has been found that attending to an expected stimulus can reverse BOLD repetition suppression (Kok et al., 2012), which may explain the selective increase of activation observed for repeats. Moreover, it has been argued that predictive coding can accommodate such findings (Auksztulewicz & Friston, 2015). In that framework, the error signal (or BOLD activation) is a result of both prediction errors and a precision weight attributed to them. Thus, increased precision of relevant item features may induce an error signal amplification as a result of even small differences between lures and repeats. Another possibility is that increased hit activation represents a change in the feedback regulation as described by Hasselmo and Schnell (1994). In this view, the activation increase for hits may result from increased shifts to a recall state. Irrespective of the exact mechanism, our data show a dissociation between LD and mnemonic discrimination. By that token, they also warrant to investigate more thoroughly whether LD is a physiologically robust index of pattern separation. (Yassa & Stark, 2011).

We did not predict activation increases to correct retrieval in HC and mPFC. However, increases in these regions have been linked to task structure learning (Kumaran et al., 2009, 2012; Kumaran & McClelland, 2012), and in the case of mPFC, to training-induced strategy learning (e.g. Kirchhoff et al., 2012). Another explanation may be a change in mPFC-mediated resolution of mnemonic interference during retrieval (see Peters et al., 2013). Peters et al. found that mPFC inactivation in rats impaired learning on an odor discrimination task and led to greater interference of learned discriminations when having to relearn item-response associations, in particular when several items had to be kept in memory. Similarly, in the present training participants faced constant interference from previous blocks whereby the same stimuli would appear in differing roles. Thus, mPFC may have been particularly involved during training, which could have led to the observed activation changes.

It should be noted that activation to correct retrieval in HC and mPFC was below baseline before training (fig. 4) and with the exception of CA3, stayed below baseline despite training-induced increases. This may put into question these regions’ involvement in the task. Accordingly, in Kelly and Garavan’s (2005) framework relevant fMRI-activation increases need to exhibit above-baseline activation. However, HC in particular is known to display a complex BOLD pattern, where negative activity can in fact be accompanied by increased neural activity (Schridde et al., 2008). Thus, a region can be involved in stimulus discrimination despite below-baseline BOLD activity (Bressler et al., 2007). We would therefore argue that the robust, training-induced increase in retrieval activation could well reflect a change in the regions’ task involvement despite below-baseline activation.

The finding of increased retrieval activation following training runs counter to the hypothesis of greater neural efficiency as an explanation for improved MD performance (see Kelly & Garavan, 2005; Poldrack, 2000). It should be noted, however, that task-fMRI activation increases following cognitive training are not uncommon (Hempel et al., 2004; Kirchhoff et al., 2012; Kühn et al., 2013; Nyberg et al., 2003; Olesen et al., 2004), even though decreases have been reported as well (Brehmer et al., 2011; Flegal et al., 2019; Miró-Padilla et al., 2019). Moreover, activation increases do not per se discard the possibility of more efficient processing, as activation increase may only be temporary. Indeed, training-induced BOLD activation changes have been reported to follow a U-shape pattern both for cortical (Hempel et al., 2004) and subcortical regions (Kühn et al., 2013). Such a pattern would suggest a form of reorganization indicated by initial activation increases, followed by increased neural efficiency reflected by activation decreases with longer training (Baykara et al., 2021). Kelly & Garavan (2005) suggest that activation increases may spill over from trainings in which task performance is not yet asymptotic. In the present training, the task remained challenging up till the end (supp. fig. 4B). A longer training paradigm with asymptotic performance and multiple measurements may shed further light on the nature and time course of the observed activation increases.

Our findings indicate that adaptivity may be crucial in inducing MD training gains and transfer. Surprisingly, we found that MD training gains were linked to a decrease rather than an increase in LD, a supposed proxy for pattern separation, and that training generally induced retrieval activation increases. This suggests that improvement in performance was reached via increased use of neural resources for memory. Finally, we found that performance improvements showed a domain-specific pattern of activation in sensory cortex, and were not induced by domain-general indices of hippocampal pattern separation. This suggests that MD training changed the domain-specific representation of stimuli rather than memory processes. Our results serve as a foundation for future research into the mechanisms that allow humans to preserve and possibly improve MD, a faculty that is known to decline with age.

## Methods

### Study 1

#### Participants

64 participants young participants (18-30 years). None reported a previous psychiatric or neurological diagnosis. All participants gave informed consent for their participation and received monetary compensation of (60€). In total, 7 participants dropped out during the course of the training, leaving a remaining sample of N = 57 (age: m = 21.56 years, sd = 9.55, 86% female). They were pseudorandomly assigned to either of three conditions, non-adaptive training (N = 19, m = 21.58 years, sd = 11.37, 73.7% female), adaptive training (N = 20, m = 23.7 years, sd = 3.61, 95% female) or control (N = 18, m = 30.83 years, sd = 15.29, 88.9% female). The study was approved by the ethics committee of the University of Magdeburg. (Probandenalter korrigieren in Datenbank und sample stats neu generieren)

#### Experimental Design

Participants performed a 2-week computerized MD task training on a newly programmed online platform (http://iknd-games.ovgu.de/MemTrain/), comprising 6 training sessions in total. Each training session lasted 45 minutes, in addition to breaks between training runs. After each training session, participants were asked to complete a feedback questionnaire that was designed to evaluate the effort, motivation and perceived difficulty evoked by the training.

All three groups completed pre- and post-test sessions, including a 6-back version of the Object-Scene task (Berron et al., 2018) and further tests of near and far memory transfer (see fig.7 for a schematic of the study outline). Near memory transfer to another MD task was assessed by the MST (Stark et al., 2013). Another form of near transfer was assessed via a task designed to primarily target retrieval rather than encoding mechanisms, namely the Object-in-Scene task (OiS). This task is supposed to tax mnemonic pattern completion, and was considered to be a near transfer task, as pattern completion also relies on hippocampal memory mechanisms, and the task uses stimuli very similar to the training stimulus set.

**Figure 7.**
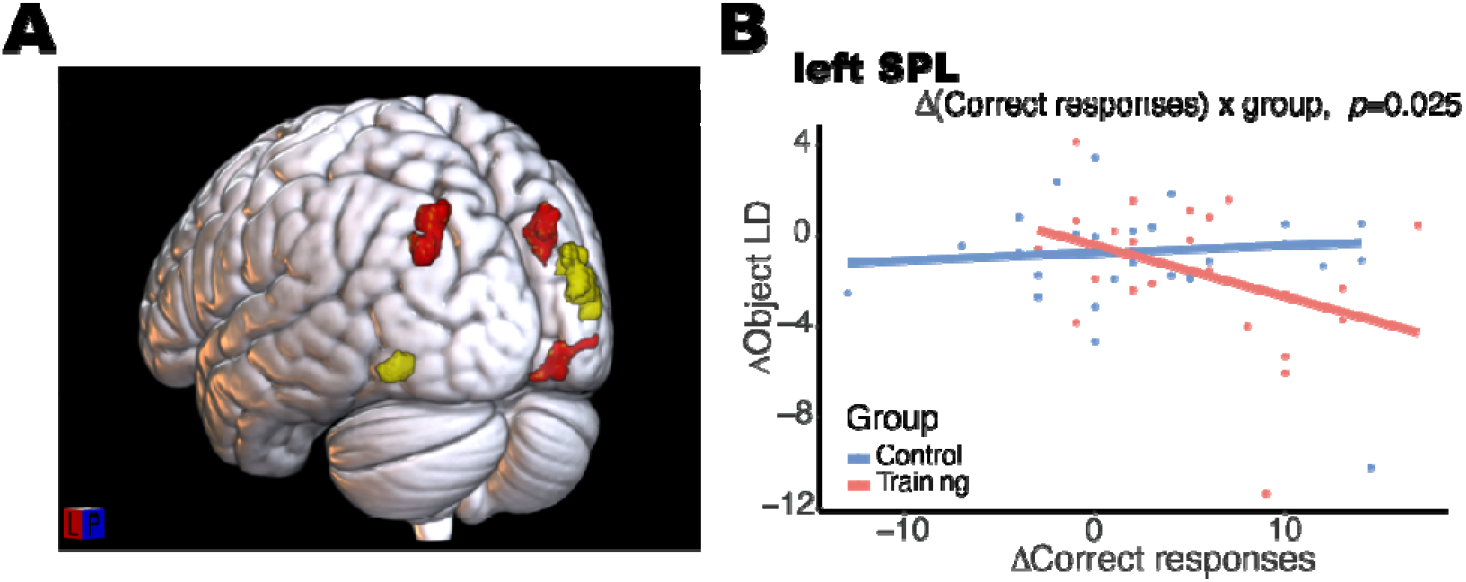
Activation changes are tied to improved OiS performance. (**A**) Change in object LD was linked to correct responses differently between groups across left superior parietal lobe, right angular gyrus and right inferior temporal gyrus (regions shown in red). These clusters had shown a link between decrease in object LD and object FAR improvements in the training group (figure 6A). Two of the clusters showed no significant interaction (shown in yellow). (**B**) The interaction effect was similar across clusters, with decreased object LD being linked to better OiS performance in the training group. Effect shown here for left SPL.

The computerized experiments outside the scanner were run on one of two screens: a Dell Optiplex9020 with integrated 23 inch screen and a resolution of 1920 * 1080, or a 21.5 inch HP ZR2240w screen with a 1920 * 1080 resolution. But for one exception due to availability, all subjects were tested on the same screen in the pre- and post-session, in order to control for potential effects of screen size. Except for the MST, which uses custom software, the tasks were run using the NBS Presentation software (version 18.1, Neurobehavioral Systems, Inc., Berkeley, CA).

#### Training Task

There were two active training groups (non-adaptive, adaptive) and one control group. Both active training groups performed an object-scene memory discrimination task, as implemented in Berron et al. (2018), notably with an extended 6-back span. The span was modified to boost difficulty and keep the task challenging, in view of participants’ young age and the expected increasing proficiency with the task. The stimuli for the training were taken from the large stimulus pool assessed in Güsten et al. (2021). This allowed us to create levels with increasing difficulty, as described further below. The training task consisted of multiple runs, each made up of 4 blocks (2 object and 2 scene blocks).

Each block contained a presentation and test trial. First, 6 stimuli were presented for 3 seconds each, separated by a short blank page (1.5 seconds). Then, a countdown (3 seconds) indicated the test trial, in which participants were presented versions of the previous stimuli, and had to indicate whether the stimulus was a repeat or a lure (max. 4.5 seconds). At each level the respective set of stimulus pairs was fixed, and the presentation of each stimulus version (version 1 or 2) was randomized for both the presentation and test trial. Participants could play up to 45 levels, and gain access to the next level upon reaching a threshold of correct responses (see supplementary material: level criterion). While object and scene blocks were presented within the same session, level statistics were calculated separately for each domain. After each run, participants could see a result page (30 seconds) containing graphs of their performance in the current and previous runs, plotted against the criterion to reach the next level. Following an optional break (max. 40 seconds), the next run started. A training session ended, when participants had spent 45 minutes in the training runs. Participants were either in the non-adaptive or adaptive training group. In the non-adaptive training, level difficulty remained constant over all levels. In the adaptive training, level difficulty increased in an approximately linear fashion (fig. 2D).

#### Control Training Task

The control group trained on a task in which they had to kill little moving neurons by clicking on them. In order to keep the visual stimulation similar to the actual training task, the background images on which the neurons moved were pseudorandomly taken from the stimulus set of adaptive training group. Task difficulty was subsequently increased in several ways: 1) reducing the presentation time of the neurons, 2) increasing the amount of neurons to eliminate, 3) increasing the speed with which the neurons move, 4) making neurons change direction and 5) introducing ’bad’ neurons appearing in red, which participants had to avoid clicking on. In order to keep the task structure close to the training task, the same general trial structure was applied, meaning that one training run consisted of 4 blocks with object and scene stimuli appearing in the background. Access to a higher level was reached by eliminating a certain percentage of neurons within a run, going from 80% until level 19 to 100% in level 34 an higher. As for the training task, once a run was finished a result page (30 seconds) appeared with information on performance and level criteria, and short breaks could be taken between runs (max. 40 seconds). Again, the daily session would end after 45 minutes spent in the training runs.

#### Transfer Tasks

As a measure of near transfer, we investigated whether performance on the original 2-back task (Berron et al., 2018) increases as a function of the 6-back training, and furthermore, whether the training regime (non-adaptive vs. adaptive) plays a role in performance change. Participants were given standardized verbal and written instructions, and then performed a 2- minute training in order to familiarize with the task. A task sequence consisted in presenting first 2 new stimuli, followed by 2 stimuli that could be either repeats or very similar lures. Stimuli were constantly displayed for 3 seconds, and fixation stars were shown between presentations, with a jittered ISI averaging 1.63 seconds and varying between 0.6 seconds and 4.2 seconds. Intervals between sequences were also jittered, but longer on average (m = 2.4s) in order to mark the end of a sequence. Participants had to report first and lure as „new“, and repeat trials as „old“ by pressing one of two buttons. A sequence exclusively contained object or scene stimuli, and the occurence of repeats and lures was counterbalanced for both domains. Moreover, the presentation of lures or repeats was not predictable for the subjects as it followed previously randomized sequences which varied for each subject and test session. A total of 56 object and 56 scene pairs were tested, with 28 first-lure and 28 first-repeat combinations for each domain. Stimulus pairs as well as their function as repeat or lure were identical for all participants. Stimulus sets for pre- and post-session differed and did also not overlap with stimuli used during the web-based training. They had furthermore been matched in difficulty based on previous evaluations.

As an alternative task testing MD, we applied the Mnemonic Similarity Task (MST Version 0,9; (Stark et al., 2013). The task consists of two phases, an encoding and a retrieval phase. Throughout the task, stimuli are presented for a duration of 2 seconds with an ISI of 0.5 seconds. In the encoding phase, a total of 128 object stimuli were presented to participants, and participants were asked to make indoor-outdoor judgments. In the retrieval phase, a total of 192 stimuli were presented, consisting of 64 repeats, 64 lures that were similar to the encoded objects, and 64 foils that were not similar to encoded stimuli. Participants could press one of three buttons, indicating whether a repeat, lure or foil had been presented. Stimulus test sets C and D were used for the pre and post-tests and their order was counterbalanced across participants. Moreover, randomized trial sequence were pseudorandomly assigned to participants.

In order to target retrieval-based learning involving object and scene stimuli, we applied the OiS developed by Xenia Grande in our lab. Note that this is a near transfer task, as it involves similar stimuli to the original task, and particularly taps on hippocampus-based memory processes. The task consists in memorizing rooms containing objects, and the specific locations of objects within a room. It was divided into an encoding and a retrieval phase. In the encoding phase, 75 trials consisted of picture presentation followed by immediate retrieval to ensure that correct encoding took place. A trial started with a random jitter (ISI: m=1s, range: 0.760-1.560) followed by the presentation of a picture that participants were asked to encode (8s). They were instructed to memorize the room, its objects and their location by imagining to mentally interact with them. After a 0.1s mask, an immediate test followed in which the same room was presented but without containing objects. A red circle on the picture acted as a cue, and participants had to indicate which object had been located in the position pointed to. To do so, they had to choose one of three possible objects that were displayed in pseudorandom order on the bottom of the screen: the correct object, the other object previously present in the picture but in another location (internal lure) as well as an object not previously present in the room at all (external lure). The choice of which object was cued was pseudorandom, and participants had to respond by pressing the left (left object), down (center object) or right (right object) arrow key. Participants had up to 3 seconds to respond, after which the original picture containing the objects was displayed again as feedback (1.5s). Once the encoding phase was over, the recall phase started immediately. The trial structure was analogous to the encoding phase, but contained no encoding part. Furthermore, a room and the location cue were shown previous to adding the three response options. Given that response time was short once the alternatives were added, participants were asked to actively recollect beforehands which object had been present in the cued location. Both the encoding and the retrieval phase contained one self-paced break. Before starting, participants did a short practice in order to familiarize with the task.

#### Statistical Analysis

Of the 57 participants that completed the training, 4 completed less than the requested 6 training sessions (m = 4.75, range: 4-5), and one completed more (7). Nevertheless, data from these participants was included in subsequent analyses. In order to estimate the effect of training on the different behavioral outcome measures, a linear mixed-effects model (LME) model was estimated for each outcome measure separately, using the lme4 package in R (Bates et al., 2015) and setting random intercepts for subjects. In addition to the main effects of group and session, we included the group x session interaction, which would suggest a training-induced change in performance. Effects of interest are then reported by applying a Type III Analysis of Variance (ANOVA) using Satterthwaite’s method to the fitted LME model. In order to further elucidate results, follow-up analyses were performed where necessary. In particular, pairwise contrasts on the effect of session were obtained from the LME for each group separately. The analyses were performed using the emmeans package, applying the kenward-roger method for the estimation of degrees of freedom.

#### Transfer Tasks

For the 2-back object-scene, we calculated the established measure of discrimination performance Pr (e.g. Berron et al., 2018) as well as HR and FAR separately for object and scene data. Due to technical error, 13 data sets only contained responses from a single run. These were nevertheless included in the analysis. The training effect for PR, HR and FAR separated by domain was estimated using a linear mixed-effects model (LME) model with the lme4 package in R (reference), and setting random intercepts for subjects. Below chance Pr scores were removed (N=1). In addition to the main effects of group and session, we included the group x session interaction, which would suggest a training-induced change in performance. Effects of interest are then reported by applying a Type III Analysis of Variance (ANOVA) using Satterthwaite’s method to the fitted LME model. Moreover, we were interested whether the training effect differed specifically between the two training groups. We therefore additionally estimated the above interaction model on the training group data only. In particular, pairwise contrasts on the effect of session were obtained from the LME for each group separately using the emmeans package (reference), and pplying the kenward-roger method for the estimation of degrees of freedom. The analyzed sample consisted of N = 53 participants (non-adaptive training: N = 18, adaptive training: N = 20, control: N = 17). For the remaining transfer tasks, the MST and the OiS, data from study 1 and 2 were pooled (see study 2 below for statistics and sample information).

#### Questionnaire Feedback

The following feedback measures were obtained and entered into analysis: excitement, fun, perceived difficulty, effort invested. We were interested in changes over training duration, and therefore specified training day as continuous variable. We therefore performed a type III Analysis of covariance (ANCOVA), using kenward-rodgers degrees of freedom with the car package in R. Regarding the analysis of training day x group interactions, given that training day is a continuous variable, pairwise contrasts could not be evaluated. We therefore directly examined the separate model interaction terms using Satterthwaite’s method for t-values. Notice that all model terms were coded relative to the adaptive training group.

### Study 2

#### Participants

60 young participants (18-30 years) were recruited at the University of Magdeburg. None reported a previous psychiatric or neurological diagnosis. All participants gave informed consent for their participation and received monetary compensation of (100€, additional prize for best players). 5 participants dropped out, so the study sample consisted of N = 55 was participants (age: m = 23.67 years, sd = 3.38, 61.8% female). Participants were pseudorandomly assigned to two groups, adaptive training (N = 28, m = 24.07, sd = 3.34, 67.9% female) or control (N = 27, m = 23.26, sd = 3.44, 55.6% female). The study was approved by the ethics committee of the University of Magdeburg.

#### Experimental Paradigm

##### Near Transfer Tasks

For the fmri task, participants received standardized verbal and written instructions, followed by 5 minutes of training the task outside of the scanner. Previous to testing, participants underwent a standardized visual screening procedure as well as a visual discrimination test involving stimuli analogous to those used in the task. If necessary, vision was corrected using MR-compatible devices. The fmri task was an adapted version of the object-scene task used in study 1: in order to make the task analogous to the one practiced on the web platform during training, the original presentation span was increased by changing the sequence length from 4 stimuli to 12 stimuli. Thus, a sequence consisted in presenting first 6 new stimuli, followed by 6 stimuli that could be either repeats or very similar lures. Stimulus duration and ISI were equivalent to study 1. The amount of tested pairs was slightly increased compared to study 1, as it had to be a multiple of the sequence length (6). Thus, a total of 60 object and 60 scene pairs were tested, with 30 first-lure and 30 first-repeat combinations for each domain, distributed over 20 sequences containing 6 stimulus pairs (12 presentations) each. The stimulus pairs used in the pre- and post-session were matched in difficulty for each combination of domain x condition combination (object repeat, object lure, scene repeat, scene lure) based on previous stimulus evaluations (Güsten et al., 2021), and did not overlap with stimuli shown during the web-based training. The MST and OiS were the same as in study 1.

##### Far Transfer Tasks

In order to test visual memory, visuospatial constructional ability and recognition ability, we applied the Rey-Osterrieth Complex Figure (ROCF, Osterrieth, 1944). The test consisted of three phases. First, participants were presented with the figure and asked to copy it as correctly as possible on a second sheet of paper. Then, after 3 minutes, a free recall followed in which participants were asked to draw the figure again from memory. A final free recall was performed ca. 30 minutes after the initial presentation of the figure. For the post-test, the Modified Taylor Complex Figure (Hubley, 1996) was used as a corresponding test in order avoid repetition effects.

In order to target verbal long-term memory as well as memory interference, we applied a modified version of verbal learning and memory test (VLMT; Helmstaedter et al., 2001). The word list for the Pre- and Posttest consisted of the A and C list of the VLMT, respectively. In order to make the task more challenging to young participants, the lists were extended to a length of 30 words by adding the lists from the CVLT. The 15-word interference list from the VLMT was kept unchanged. As an interference list for the post session, the D word list from the VLMT was used. The test procedure was as follows: first, for 5 times in a row (L 1-5), the word list was read aloud by the experimenter, followed by immediate free recall by the participant. After that, the interference list was read by the experimenter and free recall for the interference list was tested subsequently (I). Then, the participant was asked to freely recall the original word list (L 6). Finally, after ca. 30 minutes of delay from starting the test, another free recall of the word list was tested (L 7).

In order measure changes in the inhibition of cognitive interference, participants took the Stroop Task (Stroop, 1935), as implemented in the Cognitive Psychology Experiments I package (Takeuchi, 2009). The task consists in both a color naming and as well as the opposite color word naming condition.

##### Statistical Analysis - Behavioral Tasks

From the 55 participants that completed the training, 6 completed less than the requested 6 training sessions (m = 4.33, range: 3-5), and three completed more (m = 7.66, range: 7-9), but all data were still analyzed. In order to estimate the effect of training on the different behavioral outcome measures, a linear mixed-effects model (LME) model was estimated for each outcome measure separately, using the lme4 package in R (reference), and setting random intercepts for subjects. In addition to the main effects of group and session, we included the group x session interaction, which would suggest a training-induced change in performance. Effects of interest are then reported by applying a Type III Analysis of Variance (ANOVA) using Satterthwaite’s method to the fitted LME model. Pairwise contrasts on the effect of session were obtained from the LME for each group separately. The analyses were performed using the emmeans package (reference), applying the kenward-roger method for the estimation of degrees of freedom. In what follows, we describe task-specific outcome measures as well as analysis-specific exclusion information.

##### Near Transfer Tasks

For the 6-back object-scene task data, as with the 2-back task in study 1, Pr outcome measures were calculated for the object and scene domain separately.

Regarding the MST, the LDI (Lure Discrimination Index; Stark et al., 2013) was obtained, which is aimed at measuring the ability to detect lures, while controlling for response bias. As suggested by Loiotile & Courtney (2015), we furthermore extracted Pr estimates ("old"|"old" - "old"|"lure") to obtain a bias-corrected measure of discrimination.

Regarding the OiS, three different metrics of performance were analysed: we evaluated the sum of correct responses and the two error types (internal, external). Calculation of correct responses, external and internal errors was restricted to stimuli that had previously correctly been encoded. Data for the MST and OiS task from the two studies were pooled, which is why we distinguish between adaptive and non-adaptive training group. In total, 8 MST datasets could not be used due to technical error. Finally, data from participants (N = 2) that did not press the “similar” (N = 1), old” (N = 1) or "new" (N = 0) button at least once was excluded. The sample consisted of N = 102 participants (adaptive: N = 43, non-adaptive: N = 17, control: N = 42).

Regarding the OiS, 3 datasets could not be used for technical error, resulting in a final sample of N = 109 participants (adaptive: N = 46, non-adaptive: N = 19, control: N = 44).

##### Far Transfer Tasks

For both the Rey-Osterrieth Complex Figure and the Taylor Complex Figure, immediate copy (IC), 3-minute delayed recall (SD), and 30-min delayed recall (LD) were evaluated using the common 36-points measure (Meyers & Meyers, K., 1995). Data of 2 participants could not be used, leaving a sample of 53 participants (training: N = 27, control: N = 26).

For the verbal learning task 3 scores were calculated: 1) the sum of recalled items over the initial 5 word list repetitions, 2) recall after interference, consisting of the amount of recalled words before interference minus the amount recalled after (L5-L6), and 3) recall after delay, consisting of the difference between the last immediate recall and the final recall after 30 minutes (L5 - L7). Data from 5 participants could not be used due to either technical error (N = 4) or because the subject misunderstood the instructions for part of the task (N = 1), leaving a sample of 50 participants (training: N = 27, control: N = 23).

Stroop effect scores, consisting of reaction time of incongruent minus congruent trials, were calculated separately for each condition (Color name and Ink). 3 datasets could not be used due to technical error. The final sample consisted of 52 subjects (training: N = 27, control: N = 25).

#### Statistical Analysis - fMRI

##### Imaging data acquisition

All imaging data were acquired on a 3T MAGNETOM Skyra scanner (Siemens, Erlangen, Germany) using the syngo MR E11 software and a 64-channel head coil. Prior to fmri, structural T1 scans were acquired with a vendor-provided whole-head 3D magnetization prepared rapid acquisition gradient echo sequence (3D-MPRAGE) with 1mm isotropic resolution, FOV = 256 x 256mm², TR/TE/TI = 2500/4.37/1100ms, FA = 7° and BW = 140Hz/Px. Whole-brain fMRI scans were acquired in two separate runs using a 2D simultaneous multi-slice echo planar imaging sequence (SMS-Epi) developed at the Center for Magnetic Resonance Research, University of Minnesota (CMRR; Moeller et al., 2010), and the following acquisition parameters: 2 x 2mm² in-plane resolution, FOV = 212 x 212mm², TR/TE = 2200/30ms, 10% slice gap, multiband acceleration factor 2, GRAPPA 2, phase encoding (PE) direction P > A, and 2mm slice thickness. In addition, phase maps were acquired with the following parameters: TR/TE1/TE2 = 675/4.92/7.38 ms, spatial resolution = 3 mm3, and 48 slices

##### Preprocessing

Preprocessing was performed using the Statistical Parametric Mapping software (SPM, Version 12; Wellcome Trust Centre for Neuroimaging, London, UK). The first 6 scans of each run were excluded, in order to minimize signal decay artifacts. First, voxel displacement maps were calculated based on the phase maps using the FieldMap toolbox. For each run, functional images were then realigned to the first image, and unwarped using the voxel displacement maps. Coregistration matrices from individual realigned and unwarped echo-planar images (EPI) to T1-weighted images were calculated using FSL’s epi_reg function (Jenkinson & Smith, 2001). We subsequently calculated inverse coregistration matrices using convertxfm from FSL. The EPI images were then smoothed using a full-width half maximum, 4mm isotropic gaussian kernel in order to improve the signal-to-noise ratio.

##### 1st level model

A high-pass filter of 128 seconds was applied to remove low frequency signals, and a first-order autoregressive model was used to estimate temporal autocorrelation. Delta functions based on stimulus onsets were convolved with a hemodynamic response function (HRF) in order to model the regressors of the Generalized Linear Model (GLM). All task conditions were introduced as separate regressors, leading to a total of 9 regressors (first presentation, repeat trial, correct lure trial, incorrect lure trial for objects and scenes, in addition to a baseline trial consisting of noise images). One subject did not show any false alarms in the second session, and in this case only the remaining conditions were modeled. Data from the two runs were joined using spm_fmri_concatenate.m in SPM12, and run regressors were added. Additionally, the 6 motion regressors were included as covariates of no interest, in order to reduce effects of task-correlated motion (Johnstone et al., 2006). Model estimation was restricted using individual explicit gray matter masks. To obtain these, we applied a 10% threshold to the gray matter tissue probability maps obtained from SPM12’s New Segment function. We then coregistered these T1 scans to individual EPI space using FSL’s FLIRT and the inverse matrices obtained earlier. Finally, the masks were smoothed using the above mentioned 4mm kernel, in order to match the smoothing applied to the functional scans. For further analysis, we calculated LD contrasts (correct rejection (CR) minus hit) for objects and scenes separately, as well as the combined activity (average across domains; see Berron et al., 2018). In addition, we calculated contrasts for the trial types first, CR and hit by contrasting them with the noise baseline trial, both within and across domains. For the voxelwise analysis, we also obtained contrasts for domain interaction in LD ([object lure correct - object repeat correct] - [scene lure correct - scene repeat correct]). All individual contrast maps from both sessions were coregistered to T1 space using FLIRT, and subsequently normalized to MNI space using geodesic shooting (Ashburner, 2007).

##### ROI selection

We first performed a-priori ROI analyses on MTL regions, as these are consistently seen to be involved in MD tasks. We used the Penn ABC-3T ASHS MTL Atlas for T2-weighted MRI (Berron’17 protocol) from ASHS (Yushkevich et al., 2014), extracting the following regions: dentate gyrus (DG), CA3, CA1, subiculum (Sub), entorhinal cortex (ErC), perirhinal cortex (PrC; combining A35 and A36) and parahippocampal cortex (PhC). All ROI analyses were performed on unsmoothed contrast images in native space, following segmentation in ASHS. Voxels showing signal dropout were ignored when calculating mean intensity values across a ROI. For session 1, data from one subject, and for session 2, data from 2 subjects had to be excluded due to technical error in the collection of T2-weighted scans.

### 2nd level model

#### ROI analysis

All A-priori ROI analyses were carried out in R. In a first step, we excluded extreme outliers for each area and contrast, using the *identify_outliers* function from the *rstatix* package with default settings. This procedure excludes data above 1.5 x IQR above the 3^rd^ quartile or below the 1^st^ quartile. If a data point was deemed to be extreme, all contrast images for that given subject, region and session were removed. This resulted in the removal of 1.19% of the data. We then started by investigating effects at baseline (only session 1) across groups. First, we explored domain-preference of each ROI, by averaging ROI intensities across conditions, for each subject, region and domain (supp. Fig. 4). Domain-preference was then calculated as object minus scene for each subject and region, and a one-sample t-test subsequently performed for each region. We then tested repetition suppression (RS) across all ROIs, since our effect of interest, LD, is thought to be based on this phenomenon. To do so, we regressed contrast (first, hit) on the mean ROI signal together fix a fixed domain effect for each region, with random subject intercepts, in a linear mixed model using the lmer function from *lme4*. We then also evaluated in an interaction model (contrast x domain) whether the repetition sensitivity effect depends on the stimulus domain, which may suggest relevance of a region to the mnemonic processing of a particular stimulus domain. After this step, we finally explored which ROIs exhibit LD signals, using the same linear mixed models, but with CR and hits as the condition levels.

The main aim was to identify ROIs for which LD or correct retrieval activity changed through training. For this change analysis, we only included subjects for which data from both sessions was available (training = 27, control = 26). Correct retrieval activity was modelled as the average of hit, CR and FA trials ([hit + CR] / 2). For each ROI separately, we estimated an LME model with session and group and their interaction as predictors, in addition to domain as a fixed effect and a random subject slope. Moreover, we investigated domain-specific change of LD and correct retrieval. This was done within the same model framework, but including the three-way domain x group x session interaction and all lower effects, to explore whether training-related change was domain-specific. A significant three-way interaction was followed up by two-way lme models for each domain separately, to see how the group x session effect differed across domains, and finally by pairwise t-tests to investigate significant two-way interactions. For completeness, we report significant uncorrected results together with their fdr-correction.

#### Whole-brain voxelwise analysis

In addition, exploratory wholebrain voxelwise analyses were performed in SPM12 on normalized ROI activity contrasts (LD and correct retrieval) from both sessions. We again used explicit masking, obtaining a gray-matter tissue probability mask from the MNI-normalized shooting template, and thresholding it at 10% before applying 4mm isotropic gaussian smoothing. All analyses were performed on a domain-averaged LD contrast ([object lure correct - object repeat correct] + [scene lure correct - scene repeat correct])/2, and on a contrast testing a LD x domain interaction ([object lure correct - object repeat correct] - [scene lure correct - scene repeat correct]), and analogously for correct retrieval. We only report cluster-level effects that survived family-wise error rate correction (p < 0.05) at an initial cluster-defining threshold of p < 0.001. In a first step, we investigated baseline (only 1st session) ROI activity effects across both groups, in order to uncover areas showing activity consistent with LD. As we had no strong hypothesis regarding about directionality of LD and correct retrieval signal change, we calculated F-contrasts following one sample t-tests in SPM, for both the domain general as well as the interaction contrast. In a second step, we investigated activation change due to training. To do so, we first calculated change maps for both ROI activity contrasts by subtracting the contrast images of the second session from the first session. To estimate the intervention effect (group x session interaction), we ran a two-sample t-test (training vs. control group) on these change maps in SPM and used an F-contrast as we had no prior hypothesis regarding effect direction. Significant interaction effects in the change analyses were followed up by analyses performed in R on mean activity of significant clusters.

### Brain-behavior regression

#### ROI analysis

We also investigated whether individual LD activity was related to behavioral scores. Here we focused on false alarm rates, as they are closely linked to LD and were the only measure having shown significant change as a function of training. In both the a-priori ROI and the voxelwise analysis, we then focused on regions that had shown training-induced change in LD activity or correct retrieval activity (domain-general, object or scene). These were either a-priori ROIs or clusters from the voxelwise analysis that had shown a session x group interaction. In case the change effect was not domain-specific, we averaged Δ(ROI activity) and ΔFAR scores. Otherwise only the domain-specific data was used in the analysis. Mean activity was analyzed in R: we looked at individual change in the training group, setting up a linear model with ΔLD and Δ(correct retrieval) as outcome and the ΔFAR as predictor, including an intercept term, as it would otherwise affect the slope effect of interest.

A separate question was about correlation change, that is whether individual change in LD or correct retrieval was differently linked to change in performance in the two groups. This would suggest a change in the behavioral involvement of a region over training. In the a-priori ROI analysis, we calculated ΔLD and Δ(correct retrieval) scores and entered them in a linear regression with the ΔFAR x group as the effect of interest, for each domain separately as well as domain-averaged data.

#### Whole-brain voxelwise analysis

Correlation change, that is the ΔFAR x group effect was also investigated in an exploratory voxelwise analysis. To do so, change maps of LD and correct retrieval (objects, scenes and average) were entered in a flexible factorial design in SPM12, with group as factor and ΔFAR as covariate. We then estimated the group x ΔFAR interaction, obtaining ΔFAR beta estimates for each group and running F-contrasts on their difference.

#### OiS

We finally addressed whether regions of training-related change may have been responsible for OiS improvements. We therefore regressed Δ(ROI activity) on Δ(correct responses) for contrasts and ROIs having shown training-related changes in the previous analyses. Moreover, we investigated whether correlation change also emerged for the OiS. Concretely, we tested whether Δ(ROI activity) was differently related to Δ(correct responses) for training and control participants. To test this, we set up the same flexible factorial model as above, with Δ(correct responses) instead of ΔFAR as covariate.

## Acknowledgments

This work was supported by Deutsche Forschungsgemeinschaft (DFG, German Research Foundation) – Project-ID 425899996 – SFB 1436. We are grateful to Aditya Nemali for his help in setting up the training website. We thank all participants who volunteered to participate in the training. We also want to thank the staff at the IKND and the university clinic for neurology, Medical Faculty, Otto-von-Guericke University for assistance in testing and scanning the participants.

## Supplementary information

### Feedback Questionnaire Study 1

Regarding excitement, there was a weak trend for group difference, with adaptive training appearing most exciting (m = 2.96, sd = 1.04), followed by the non-adaptive training (m = 2.71, sd = 0.93) and the control training regime (m = 2.58, sd = 1.16; F(2, 106.4432) = 2.4242, p = 0.0934). On the other hand, excitement did not significantly change over training days (F(1, 254.1922) = 0.3918, p = 0.5319). Finally, there was an interaction of group by training day (F(2, 253.6046) = 3.9369, p = 0.0207). The separate group by training interaction estimates for the non-adaptive group (*t*(253.53) = -1.6734, p = 0.0955) and the control group (estimate: 0.07, *t*(254.05) = 1.1983, p = 0.2319) indicate that the interaction was driven mainly by a decrease in excitement in the non-adaptive training over training session relative to the adaptive training. The training regimes differed in terms of the fun experienced, with the adaptive (m = 3.09, sd = 1.09) and non-adaptive (m = 3.07, sd = 1.02) training groups experiencing more fun than the controls (m = 2.57, sd = 0.97; F(2, 158.5176) = 6.998, p = 0.0012). Experienced fun did not significantly change along the training (F(1, 255.2705) = 1.3035, p = 0.2546). However, there was a significant group x training day interaction (F(2, 254.7036) = 3.6938, p = 0.0262), mainly driven by a positive interaction term for the controls (estimate: 0.14, *t*(255.73) = 2.0815, p = 0.0384), suggesting that the difference in fun between control training and adaptive training reduced over the training. Importantly, the groups did not differ in the reported effort that they invested in the training (F(2, 125.9696) = 0.1583, p = 0.8538), while there was trend for change over the training (F(1, 254.6184) = 3.8051, p = 0.0522), caused by a decrease in effort over sessions (estimate = 254.21). Finally, the groups differed in terms of the perceived difficulty, with the control training being perceived most difficult (m = 3.66, sd = 0.93), followed by the adaptive training (m = 3.46, sd = 0.89) and lastly the non-adaptive training (m = 3.03, sd = 0.81; F(2, 135.4208) = 3.5544, p = 0.0313). Moreover, there was a trend of difficulty increase over the training (F(1, 254.8114) = 3.341, p = 0.0687)

### Feedback Questionnaire Study 2

We found no significant effects of group or training day for either excitement (group: F(1, 113.569) = 0.0566, p = 0.8123; training day: F(1, 210.3018) = 1.7535, p = 0.1869), fun

(group: F(1, 165.2502) = 0.46, p = 0.4986; training day: F(1, 216.2443) = 0.0998, p =

0.7524), effort invested (group: F(1, 150.752) = 1.7135, p = 0.1925; training day: F(1, 214.3602) = 0.2752, p = 0.6004), or perceived difficulty (group: F(1, 168.5614) = 0.0751, p =

0.7844; training day: F(1, 216.7128) = 2.575, p = 0.11).

---

**Supp. fig. 1.**
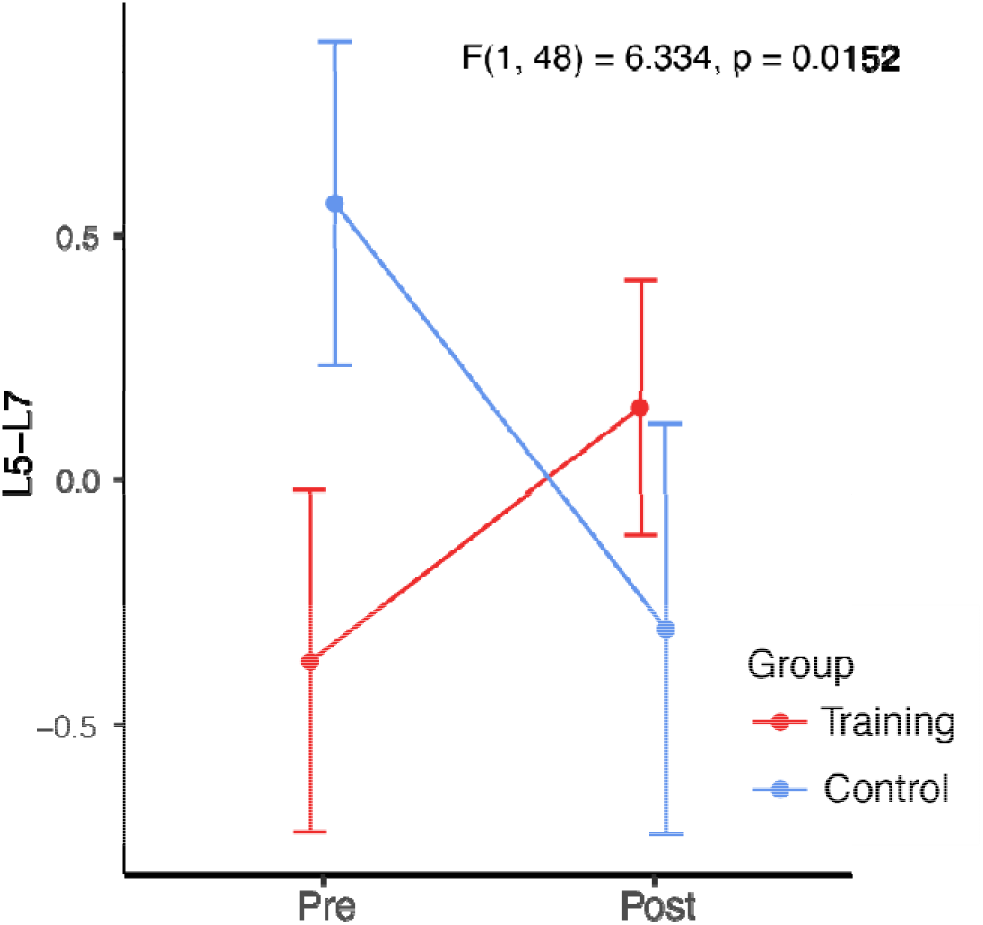
Word List Recall: Group x Session interaction data of Delayed Recall (L7) minus Direct Recall (L5). While in the pre session, the control group had worse L5 -L7 scores, the training group performed worse in the post session. This was due to an mean decrease in performance for the training group, and a mean decrease for the control group. Figure shows mean and standard error (SE).

**Supp. fig. 2.**
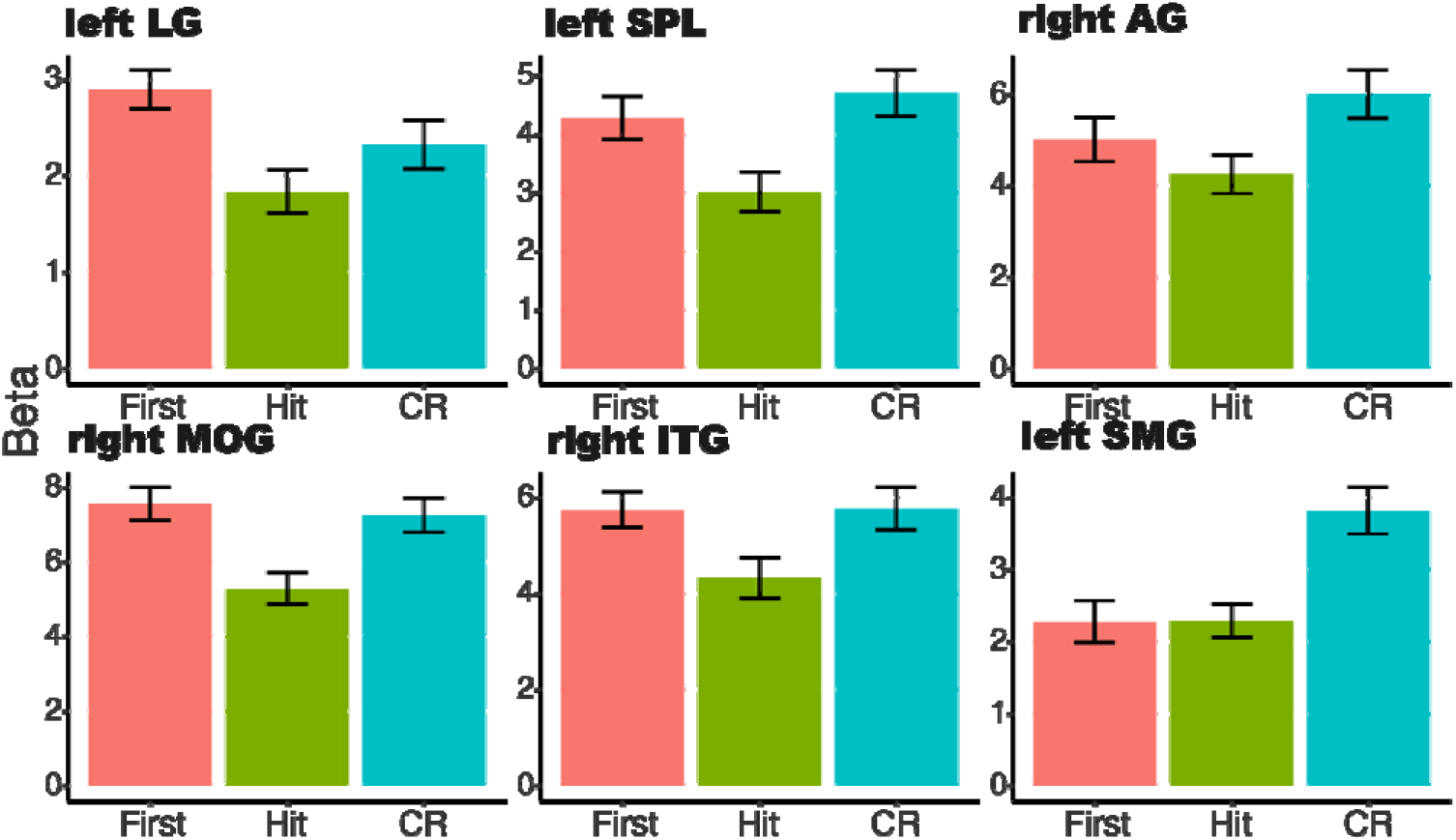
Baseline activation. Activation profiles in regions showing correlation change with ΔLD.

**Supp. fig. 3.**
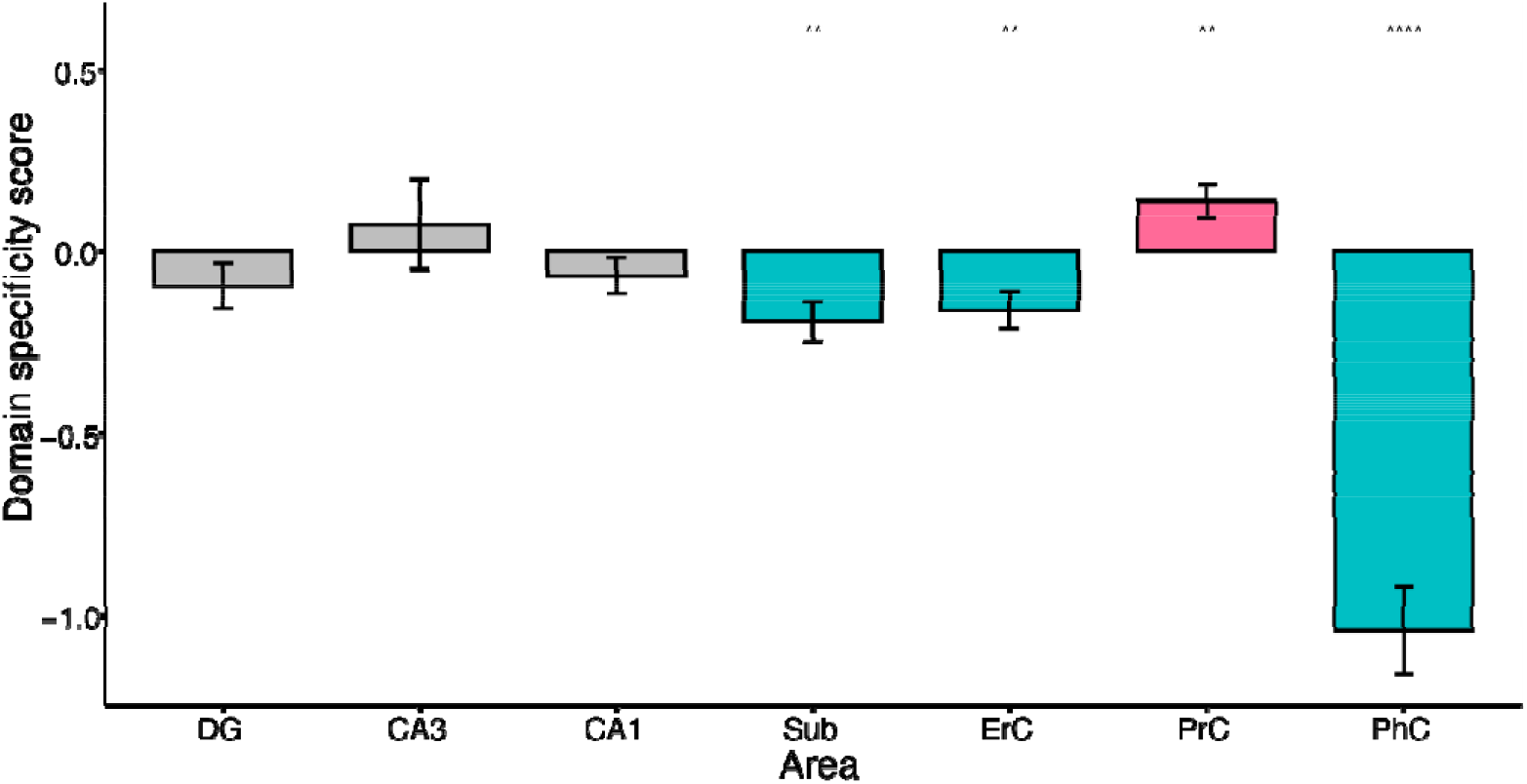
Domain-specificity of MTL regions at baseline. Red/blue = stronger overall activation to objects/ scenes.

**Supp. fig. 4.**
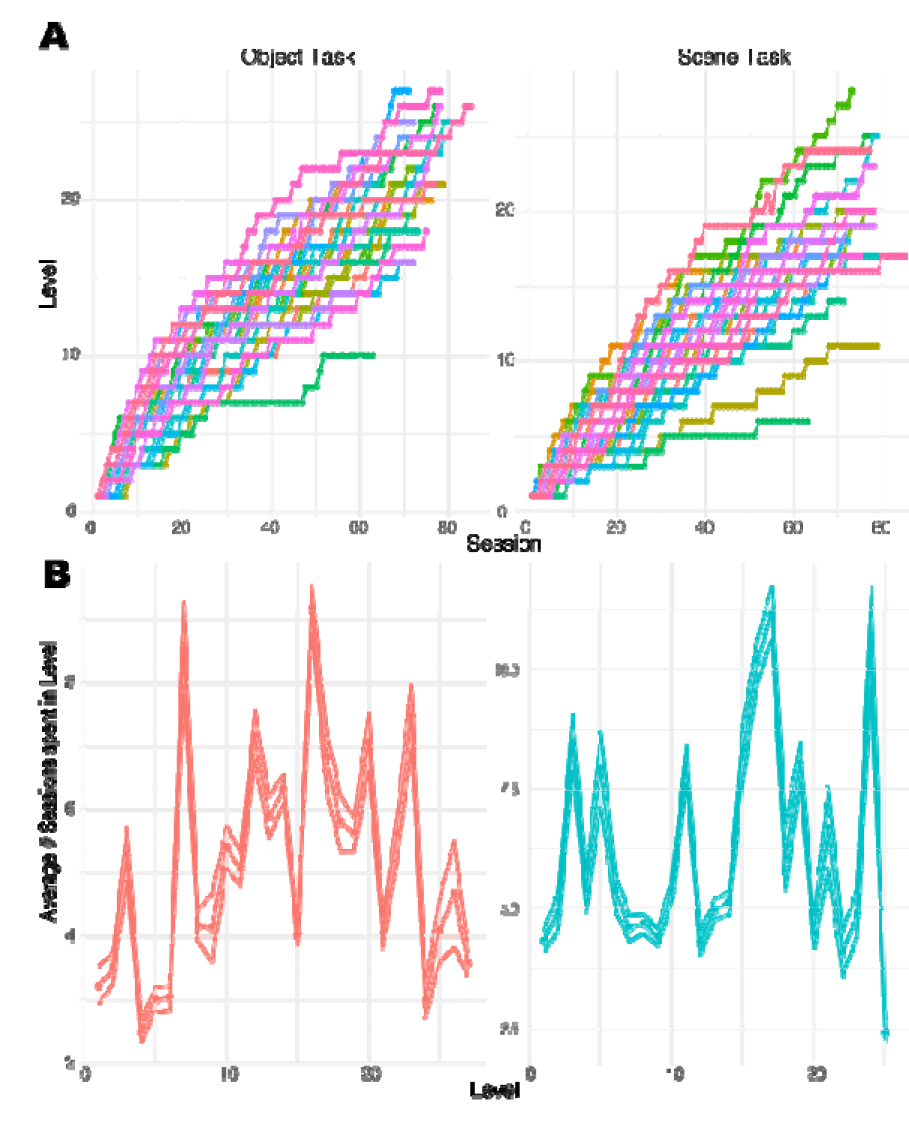
A. Level progress of the adaptive training group over the training sessions for the object and scene task. Each line describes the trajectory of one participant. No participant reached the last level (45). **B** Average number of sessions the adaptive training group spent in the object and scene levels. Performance was not asymptotic (at later levels, participants still needed more than 1 session to reach the next level). Error band shows standard error.

**Supp. table 1.** Object-Scene Task: Study 1. HR=hit rate, FAR= false alarm rate, Pr= hit rate - false alarm rate, o/s= object/ scene. p-statistic: group x session interaction.

| Adaptive | Non-Adaptive | Control |
| --- | --- | --- |

|  | Pre |  | Post |  | Pre |  | Post |  | Pre |  | Post |  |  |
| --- | --- | --- | --- | --- | --- | --- | --- | --- | --- | --- | --- | --- | --- |
| *Score* | m | SE | m | SE | m | SE | m | SE | m | SE | m | SE | p |
| *HR<sub>o</sub>* | 0.89 | 0.03 | 0.86 | 0.03 | 0.86 | 0.03 | 0.84 | 0.03 | 0.83 | 0.03 | 0.83 | 0.03 | 0.8397 |
| *FAR<sub>o</sub>* | 0.47 | 0.04 | 0.21 | 0.04 | 0.35 | 0.04 | 0.3 | 0.04 | 0.44 | 0.04 | 0.37 | 0.04 | 0.001 |
| *HR<sub>s</sub>* | 0.9 | 0.03 | 0.88 | 0.03 | 0.9 | 0.03 | 0.81 | 0.03 | 0.83 | 0.03 | 0.84 | 0.03 | 0.1737 |
| *FAR<sub>s</sub>* | 0.45 | 0.04 | 0.26 | 0.04 | 0.39 | 0.04 | 0.28 | 0.04 | 0.44 | 0.04 | 0.36 | 0.04 | 0.1054 |
| *Pr<sub>o</sub>* | 0.41 | 0.04 | 0.65 | 0.04 | 0.51 | 0.04 | 0.62 | 0.04 | 0.39 | 0.04 | 0.46 | 0.04 | 0.0292 |
| *Pr<sub>s</sub>* | 0.44 | 0.04 | 0.62 | 0.04 | 0.51 | 0.05 | 0.61 | 0.05 | 0.41 | 0.05 | 0.48 | 0.05 | 0.1223 |

**Supp. table 2.** Object-Scene Task: Study 2. p-statistic: group x session interaction.

|  | Training |  |  |  | Control |  |  |  |  |
| --- | --- | --- | --- | --- | --- | --- | --- | --- | --- |
|  | Pre |  | Post |  | Pre |  | Post |  |  |
| *Score* | m | SE | m | SE | m | SE | m | SE | p |
| *HR<sub>o</sub>* | 0.92 | 0.01 | 0.91 | 0.01 | 0.91 | 0.02 | 0.9 | 0.02 | 0.5248 |
| *FAR<sub>o</sub>* | 0.35 | 0.02 | 0.16 | 0.02 | 0.36 | 0.02 | 0.29 | 0.02 | 0.0014 |
| *H<sub>s</sub>* | 0.87 | 0.02 | 0.9 | 0.02 | 0.89 | 0.02 | 0.88 | 0.02 | 0.102 |
| *FAR<sub>s</sub>* | 0.43 | 0.02 | 0.21 | 0.02 | 0.43 | 0.02 | 0.32 | 0.02 | 0.0023 |
| *Pr<sub>o</sub>* | 0.57 | 0.03 | 0.74 | 0.03 | 0.54 | 0.03 | 0.61 | 0.03 | 0.0166 |
| *Pr<sub>s</sub>* | 0.44 | 0.03 | 0.69 | 0.03 | 0.46 | 0.03 | 0.57 | 0.03 | 0.0017 |

**Supp. table 3.** OiS: Study 1&2. p-statistic: group x session interaction.

|  | Adaptive |  |  |  | Non-Adaptive |  |  |  | Control |  |  |  |  |
| --- | --- | --- | --- | --- | --- | --- | --- | --- | --- | --- | --- | --- | --- |
|  | Pre |  | Post |  | Pre |  | Post |  | Pre |  | Post |  |  |
| *Score* | m | SE | m | SE | m | SE | m | SE | m | SE | m | SE | p |
| $\Sigma$ | 49.22 | 1.4 | 55.07 | 1.4 | 51.53 | 2.18 | 53.21 | 2.18 | 52.14 | 1.43 | 54.14 | 1.43 | 0.0126 |
| *Ext. Errors* | 7.2 | 0.59 | 6.11 | 0.59 | 5.58 | 0.91 | 6.47 | 0.91 | 5.36 | 0.6 | 5.75 | 0.6 | 0.1129 |
| *Int. Errors* | 12.52 | 0.77 | 10.89 | 0.77 | 11.37 | 1.21 | 11.84 | 1.21 | 11.16 | 0.79 | 10.2 | 0.79 | 0.2244 |

**Supp. table 4.** Verbal Learning: Study 1&2. Σ L1 -L5= sum of remembered word in list 1 -5 . L5 - L6 = change of recalled words after interference. L5- L7= change of recalled words after 30. p-statistic: group x session interaction.

|  | Training |  |  |  | Control |  |  |  |  |
| --- | --- | --- | --- | --- | --- | --- | --- | --- | --- |
|  | Pre |  | Post |  | Pre |  | Post |  |  |
| *Score* | m | SE | m | SE | m | SE | m | SE | p |
| $\Sigma L1-L5$ | 120.15 | 2.67 | 124.52 | 2.67 | 118.09 | 2.89 | 122.96 | 2.89 | 0.892 |
| *L5-L6* | 0.85 | 0.38 | 0.63 | 0.38 | 0.87 | 0.41 | -0.22 | 0.41 | 0.171 |
| *L5-L7* | -0.37 | 0.33 | 0.15 | 0.33 | 0.57 | 0.36 | -0.3 | 0.36 | 0.015 |

**Supp. table 5.** MST: Study 1&2. Pr: hit-rate - false alarm rate. LDI: lure-discrimination index. p-statistic: group x session interaction.

|  | Adaptive |  |  |  | Non-Adaptive |  |  |  | Control |  |  |  |  |
| --- | --- | --- | --- | --- | --- | --- | --- | --- | --- | --- | --- | --- | --- |
|  | Pre |  | Post |  | Pre |  | Post |  | Pre |  | Post |  |  |
| *Score* | m | SE | m | SE | m | SE | m | SE | m | SE | m | SE | p |
| *Pr* | 0.39 | 0.02 | 0.39 | 0.02 | 0.46 | 0.04 | 0.48 | 0.04 | 0.41 | 0.02 | 0.46 | 0.02 | 0.267 |
| *LDI* | 0.34 | 0.03 | 0.38 | 0.03 | 0.45 | 0.04 | 0.49 | 0.04 | 0.35 | 0.03 | 0.41 | 0.03 | 0.731 |

**Supp. table 6.** Rey-Osterrieth (ROCF) &Taylor (MTCF): Study 1&2. C: Copy, SD: 3- minute delayed recall, D: 30-minute delayed recall. p-statistic: group x session interaction.

|  | Training |  |  |  | Control |  |  |  | p |
| --- | --- | --- | --- | --- | --- | --- | --- | --- | --- |
|  | Pre |  | Post |  | Pre |  | Post |  |  |
| *Score* | m | SE | m | SE | m | SE | m | SE |  |
| *C* | 35.22 | 0.22 | 35.06 | 0.22 | 35.33 | 0.22 | 35.31 | 0.22 | 0.6924 |
| *SD* | 27.48 | 0.68 | 32.09 | 0.68 | 28.92 | 0.69 | 31.87 | 0.69 | 0.1482 |
| *D* | 27.11 | 0.71 | 31.69 | 0.71 | 28.13 | 0.72 | 31.94 | 0.72 | 0.4874 |

**Supp. table 7.** Stroop Task: Study 2. Word = target is the color name. Ink = target is the ink color of shown letters. p-statistic: group x session interaction.

|  | Training |  |  |  | Control |  |  |  | p |
| --- | --- | --- | --- | --- | --- | --- | --- | --- | --- |
|  | Pre |  | Post |  | Pre |  | Post |  |  |
| *Score* | m | SE | m | SE | m | SE | m | SE |  |
| *Word* | 55.34 | 516.85 | 71.48 | 516.85 | 45.99 | 516.85 | -1002.03 | 516.85 | 0.306 |
| *Ink* | 88.96 | 254.2 | 98.03 | 254.2 | -368.5 | 254.2 | -132.5 | 254.2 | 0.656 |

**Supp. table 8.** Effect of training (session x group) on correct retrieval.

| Contrast | Domain | Cluster | Peak | MNI (mm) |  |  | Region |  |  |
| --- | --- | --- | --- | --- | --- | --- | --- | --- | --- |
|  |  | p(FWE-corr) | size | Z | x | y | z |  |  |
| *Correct Retrieval* | General | 0.002 | 815 | 3.94 | -7 | 52 | -8 | L | MFC |
|  |  |  |  | 3.8 | 0 | 51 | -13 |  |  |
|  |  |  |  | 3.73 | -3 | 53 | 0 |  |  |

